# Dynamic mechanisms of CRISPR interference by *Escherichia coli* CRISPR-Cas3

**DOI:** 10.1101/2021.07.18.452824

**Authors:** Kazuto Yoshimi, Kohei Takeshita, Noriyuki Kodera, Satomi Shibumura, Yuko Yamauchi, Mine Omatsu, Yayoi Kunihiro, Masaki Yamamoto, Tomoji Mashimo

## Abstract

Type I CRISPR-Cas3 uses an RNA-guided multi Cas-protein complex, Cascade, which detects and degrades foreign nucleic acids via the helicase-nuclease Cas3 protein. Despite many studies using cryoEM and smFRET, the precise mechanism of Cas3-mediated cleavage and degradation of target DNA remains elusive. Here we reconstitute the CRISPR-Cas3 system *in vitro* to show how the *Escherichia coli* Cas3 (EcoCas3) with EcoCascade exhibits collateral non-specific ssDNA cleavage and target specific DNA degradation. Partial binding of EcoCascade to target DNA with tolerated mismatches within the spacer sequence, but not the PAM, elicits collateral ssDNA cleavage activity of recruited EcoCas3. Conversely, stable binding with complete R-loop formation drives EcoCas3 to nick the non-target strand (NTS) in the bound DNA. Helicase-dependent unwinding then combines with *trans* ssDNA cleavage of the target strand and repetitive *cis* cleavage of the NTS to degrade the target dsDNA substrate. High-speed atomic force microscopy demonstrates that EcoCas3 bound to EcoCascade repeatedly reels and releases the target DNA, followed by target fragmentation. Together, these results provide a revised model for collateral ssDNA cleavage and target dsDNA degradation by CRISPR-Cas3, furthering understanding of type I CRISPR priming and interference and informing future genome editing tools.

## Main

The clustered-regularly-interspaced-short-palindromic-repeats (CRISPR) CRISPR-associated proteins (Cas) system allows for adaptive immunity in prokaryotes. CRISPR protein complexes comprise two classes, with each class classified into three types where Class 1 includes type I, III, IV and Class 2 includes type II, V, and VI ^1^. Class 1 systems use multiple different Cas proteins, while Class 2 effectors contain only a single protein. To date, much attention has focused on the mechanism of Class 2 effectors, such as type II Cas9, type V Cas12, and type VI Cas13, given their practical applications in genome editing and manipulation ^2–6^}. Type 1 systems are also now emerging as tools for genome and transcriptome manipulation in microbiota ^7, 8^ and eukaryotic cells ^9–11^. Binding of type I CRISPR-Cas effectors to DNA sequences in the absence of Cas3 leads to transcriptional repression in bacteria ^7^ and human cells ^9^. For DNA editing, introduction of the Cascade multi Cas-protein complex, CRISPR RNA (crRNA), and the Cas3 helicase-nuclease into mammalian cells results in long-range chromosomal deletions in target DNA ^10, 11^. The long-range deletions generated by Cas3 contrasts with smaller deletions (or indels) generated by Cas9/Cas12 editing, and has led to the descriptions of DNA shredder and scissors, respectively.

Considerable efforts have been devoted to understanding the mechanism of CRISPR interference by type I CRISPR ^12–26^. Several cryo-electron microscopy (EM) structures of type I CRISPR complexes have been solved, revealing seahorse-shaped structures containing Cas5, Cas6, multiple Cas7, Cas8 (Cse1), which recognizes the PAM, and two Cas11 (Cse2) (Fig. 1a). Type I CRISPR systems target homologous regions of double-stranded DNA (dsDNA) for degradation through two major steps: recognition of a target DNA by Cascade complex surveillance, and cleavage of the DNA by Cas3 that is recruited by the Cascade complex ^23–25^. In the first step, the Cascade complex, including Cas8, scans the PAM (protospacer adjacent motif) and initiates DNA unwinding at the PAM ^27^. Subsequently, crRNA hybridization with the target DNA strand (TS) leads to displacement of the non-target strand (NTS), forming a three-stranded nucleic acid structure known as an R-loop ^14–16, 18, 19, 26^. Complete formation of the R-loop induces a conformational change in the Cascade complex that enables recruitment of Cas3 ^15, 19, 28^. The recruited Cas3, a protein with an SF2 (Superfamily 2) helicase domain and a HD (histidine-aspartate) nuclease domain, degrades the target DNA in a unidirectional ATP-dependent manner according to the following steps: nicking the NTS at the R-loop, loading onto the ssDNA, and unwinding the DNA while degrading the DNA ^13, 15, 25, 29, 30^. Recently, single-molecule Förster resonance energy transfer (smFRET) experiments have shown that Cas3 can remain associated with Cascade to cleave ssDNA by a reeling mechanism ^12^. However, Cas3 can also break free of Cascade and translocate on its own through the target DNA ^20^. It is unclear whether Cas3 degrades DNA during independent translocation and how Cas3 with a single HD domain can degrade both the NTS and the TS of dsDNA ^31, 32^ (Fig. 1a).

**Figure. 1:**
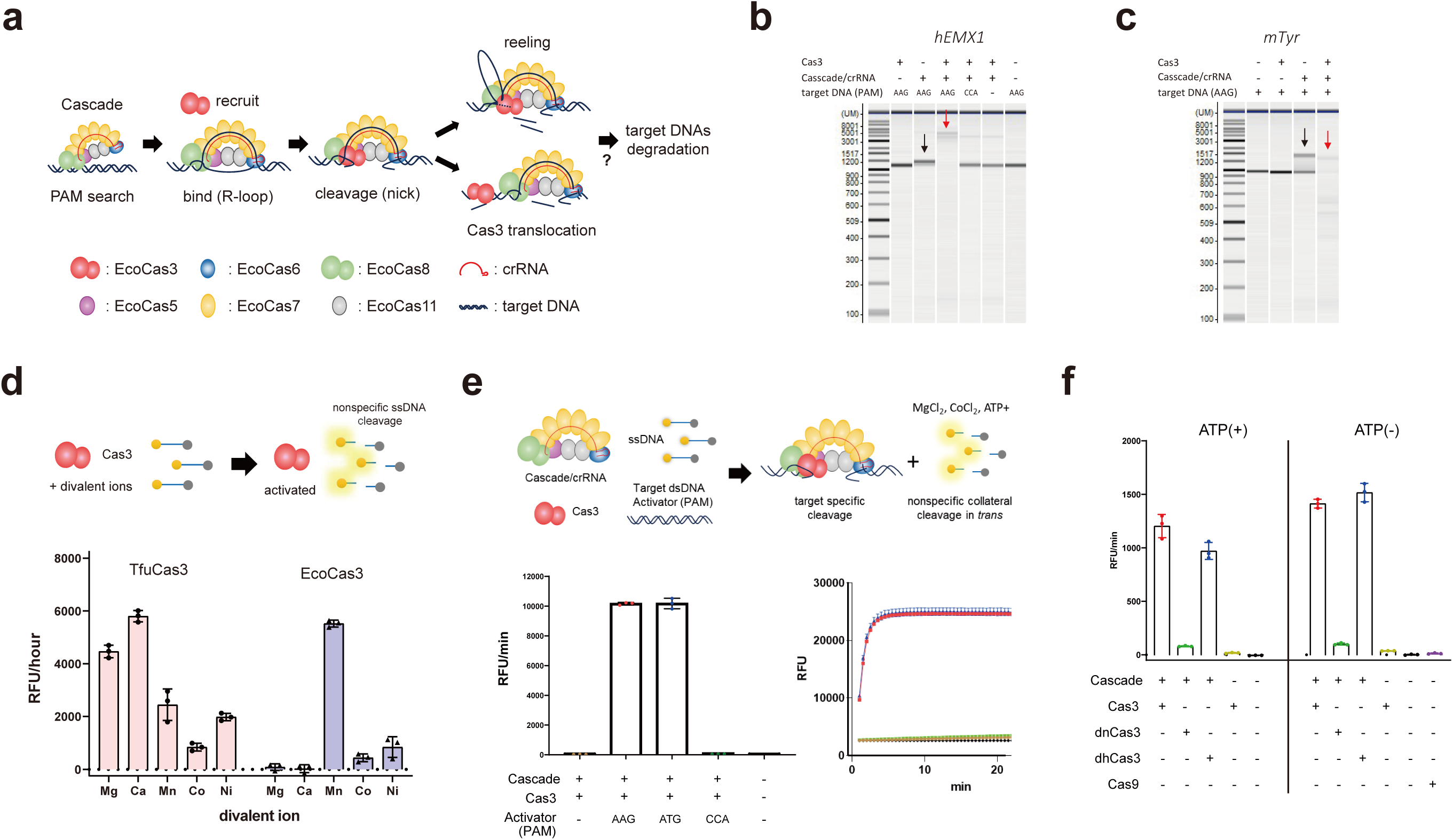
*In vitro* reconstituted EcoCas3-EcoCascade/crRNA complex cleaves nonspecific ssDNA in *trans*. (a) Schematic depiction of the known type I CRISPR interference mechanism. (b,c) Electrophoretic mobility shift assay (EMSA) and DNA degradation assay. (b) EcoCascade/crRNA complex binds to target plasmids containing *hEMX1* spacer sequences flanked by PAM (AAG) (black arrow). EcoCas3 recruited into EcoCascade degrades the target DNA (red arrow). (c) EcoCascade binds to PCR products of the target DNA (*mTyr*) (black arrow), which is degraded by EcoCas3 (red arrow). (d) Activation of EcoCas3 and TfuCas3 by divalent metal ions (Mg^2+^, Ca^2+^, Mn^2+^, Co^2+^ and Ni^2+^). Fluorescent dye-quencher (FQ)-labeled ssDNA probes measured promiscuous ssDNA cleavage activity. RFU: relative fluorescence unit. (e) Collateral ssDNA cleavage activity measured by incubation of EcoCas3-EcoCascade/crRNA complex with a 60-bp dsDNA Activator containing a target sequence flanked by a PAM and an FQ-labeled ssDNA probe in reaction buffer containing MgCl_2_, CoCl_2_, and ATP for 10 min at 37°C. Quantitatively represented by RFU per min (left) or RFU at 10 min (right). (f) EcoCas3 HD domain H74A (dead nuclease mutant, dnCas3), abolished collateral cleavage activity, while SF2 motif III S483A/T485A (dead helicase mutant, dhCas3) showed collateral cleavage activity. Collateral activity in ATP reaction buffer (+) was at the same level as that in ATP-free buffer (-) for wild-type EcoCas3 and the dhCas3 mutant. SpCas9 did not exhibit any collateral cleavage activity.

This study reveals that the *Escherichia coli* Cascade crRNA complex (EcoCascade) assembled with *E. coli* Cas3 (EcoCas3) exhibits collateral *trans*-cleavage activity on a non-specific ssDNA. We show that unstable EcoCascade binding with partial R-loop formation with EcoCas3 mediates this collateral *trans*-cleavage, but does not lead to double-stranded DNA cleavage. In contrast, stable EcoCascade binding with locked R-loop construction provides *cis* cleavage of the NTS with helicase-dependent *trans* separation and cleavage of the TS, resulting in progressive degradation of target dsDNA substrates. *In vitro* experiments using high-speed atomic force microscopy (hs-AFM) also demonstrate that EcoCas3 remains tightly associated with the EcoCascade, which repeatedly reels and releases the target DNA, followed by target degradation. These findings provide insight into the mechanism of type I CRISPR-Cas3 priming and interference against a foreign DNA.

## Results

### *In vitro* reconstitution of *Escherichia coli* CRISPR-Cas3 interference

*E. coli* CRISPR-Cas3 is generally well-characterized type I CRISPR complexes *in vitro* and *in vivo* ^31–34^. However, recombinant EcoCas3 protein is difficult to purify because of poor solubility and propensity to aggregate at 37°C ^24, 25, 29, 35^. To overcome this problem, we attempted using co-expressed HtpG chaperon ^36^ and/or low temperature growth at 20°C ^23, 24^. However, these approaches produced only a limited amount of protein that was highly aggregated (Extended data Fig. 1a). This is in contrast to isolated *Thermobifida fusca* Cas3 (TfuCas3) protein produced by the *E. coli* bacterial expression system at 37°C ^10, 15, 30^ (Extended data Fig. 1b). We then used Sf9 insect cell with a baculovirus expression system for protein expression at 20°C ^37^ (Extended data Fig. 2a). EcoCas3 protein purified from Sf9 cells was soluble and ∼95% homogeneous, as evaluated by sodium dodecyl sulfate-polyacrylamide gel electrophoresis (SDS-PAGE) (Extended data Fig. 2b). EcoCascade proteins and crRNA were co-expressed in *E. coli* JM109(DE3), purified using Ni-NTA resin, and separated by size exclusion chromatography (SEC), as previously reported ^23, 38^ (Extended data Fig. 2c). Purified EcoCascade/crRNA ribonucleoproteins (RNPs) were size-evaluated by SDS-PAGE (Extended data Fig. 2d) and were consistent with those of previous reports ^23, 38^. Various methods using Tycho NT.6 protein stability measurements (Extended data Fig. 3) and the ProteoStat protein aggregation assay (Extended data Fig. 4) indicated that the temperature-dependent stability and aggregation onset temperature of EcoCas3 was mostly consistent with a mesophilic protein ^24, 25, 29, 35^.

Next, we sought to evaluate whether co-purified recombinant EcoCascade-crRNA RNPs were competent for target DNA recognition and degradation. EcoCascade-crRNA RNPs bound to supercoiled (SC) plasmids composed of *hEMX1* spacer sequences flanked by a PAM (AAG). However, these RNPs did not bind to SC plasmids when they included spacer sequences flanked by a nonPAM (CCA) (Fig. 1b and Extended data Fig. 5a). Binding of EcoCascade-crRNA RNPs was also observed for linear dsDNA molecules harboring other target sequences (*mTyr* and *rIl2rg* genes) (Fig. 1c and Extended data Fig. 5b). Exchanging nucleotide pairs between crRNAs and target sequences abolished binding, indicating the specificity of target recognition by EcoCascade RNPs (Extended data Fig. 5c). Finally, assembly of EcoCascade RNPs with EcoCas3 specifically degraded SC plasmids (*hEMX1*) and linear dsDNA (*mTyr*) in the presence of ATP and Mg^2+^ ^23, 24, 29, 35, 39^ (Fig 1b,c). Overall, our data indicate functional reconstitution of recombinant EcoCas3-EcoCascade-cRNA complexes.

### The EcoCas3-EcoCascade-crRNA complex nonspecifically cleaves ssDNAs in *trans*

Cas3 proteins from *Streptococcus thermophiles, Methanocaldococcus jannaschii*, and *Thermus thermophilus* can exhibit indiscriminate, divalent cation-dependent ssDNase activity in the absence of Cascade ^29, 35, 39^. Using fluorescent dye-quencher (FQ)-labeled ssDNA probes, we found that EcoCas3 and TfuCas3 also exhibit nonspecific ssDNA cleavage in a metal-dependent manner, although the dependency was different between the two bacteria (Fig. 1d). TfuCas3 cleaved ssDNA with all divalent ions tested, whereas EcoCas3 was only activated with Mn^2+^ and Ni^2+^, consistent with previous results ^24^. Type V Cas12a, an RNA-guided DNase ^40^, and type VI Cas13, an RNA-guided RNase ^41^, engage in collateral cleavage of nearby non-specific nucleic acids after their targeted activity. To investigate whether Cas3 also possesses collateral ssDNA cleavage activity, we assembled EcoCas3, EcoCascade RNPs, 60-bp dsDNA fragments containing target sequences flanked by a PAM (targeted Activator), and a untargeted ssDNA^42^. We found that targeted degradation triggered untargeted degradation of both circular M13 phage ssDNA and linearized long ssDNA, but not of circular pBlueScript dsDNA (Extended data Fig. 6a). As is the case with target dsDNA degradation by EcoCascade RNPs and EcoCas3 (Fig 1b,c), this collateral ssDNA cleavage was dependent upon the presence of a PAM in the targeted nucleic acid (Extended data Fig. 6a). These results indicate that either some metal ions or Cascade target-binding by R-loop formation can induce EcoCas3-dependent non-specific ssDNA cleavage activity *in vitro*.

To quantitatively measure collateral ssDNA cleavage activity we used a FQ-labeled untargeted ssDNA probe ^40, 41^ (Fig. 1e), which is used in CRISPR-based diagnostics as a platform for rapid and sensitive nucleic acid detection, for example in Covid-19 test kits ^43–45^. Consistent with the results of the M13/linear ssDNA cleavage (Extended data Fig. 6a), EcoCas3 showed collateral ssDNA cleavage in a PAM-dependent manner (with a PAM of AAG or ATG, but not CCA) (Fig. 1e and Extended data Fig. 6b,c). Fluorescent reporter DNA oligonucleotides (DNaseAlert™ IDT) also confirmed this collateral cleavage activity (Extended data Fig. 7a), whereas fluorescent reporter RNA oligonucleotides (RNaseAlert™, IDT) detected little or no collateral RNase activity (Extended data Fig. 7b). We previously showed that mutants of EcoCas3 in the HD domain (H74A, dead nuclease, dnCas3), and SF2 motif III (S483A/T485A, dead helicase, dhCas3) abolished target DNA degradation in human cells ^11^. In the collateral cleavage assay, the dnCas3 mutant abolished all cleavage activity. This indicates that *trans* cleavage of non-specific ssDNA was catalyzed by the HD domain (Fig. 1f). In contrast, the collateral cleavage activity of the dhCas3 mutant was only slightly lower than that of wild-type EcoCas3 (Fig. 1f). In ATP-free reaction buffer (-), the collateral activity of the EcoCas3 protein was at the same level as that of wild-type EcoCas3 and the dhCas3 mutant in ATP (+) buffer (Fig. 1f). Together, these results indicate that the nuclease and helicase activities of EcoCas3 are required for target DNA degradation, but only the nuclease activity is required for collateral cleavage.

### PAM recognition is a prerequisite for collateral ssDNA cleavage by Cas3 but not Cas12a

Having determined that collateral ssDNA cleavage by the EcoCas3-EcoCascade complex is PAM-dependent (Fig. 1e), we sought to further characterize the specificity of PAM recognition by screening all 64 possible target sites containing each of the three-nucleotide PAM sequences (Fig. 2a,b and Extended data Fig. 8a,b). We observed collateral cleavage activity with 14 PAM types, with the highest activity from AAG and ATG, followed by GAG, AAA, AAC, TAG, and AGG. There was no cleavage when the first or second PAM nucleotide was C or the third nucleotide was T (Fig. 2a and Extended data Fig. 8a). This PAM recognition specificity for *trans* cleavage activity matched the results from an *in vivo* high-throughput CRISPR-interference assay ^46^. In contrast, LbaCas12a showed collateral cleavage activity with almost all 64 PAM types, with the highest activity with GGGG and the lowest with GCCG (Fig. 2b and Extended data Fig. 8b).

**Figure 2.**
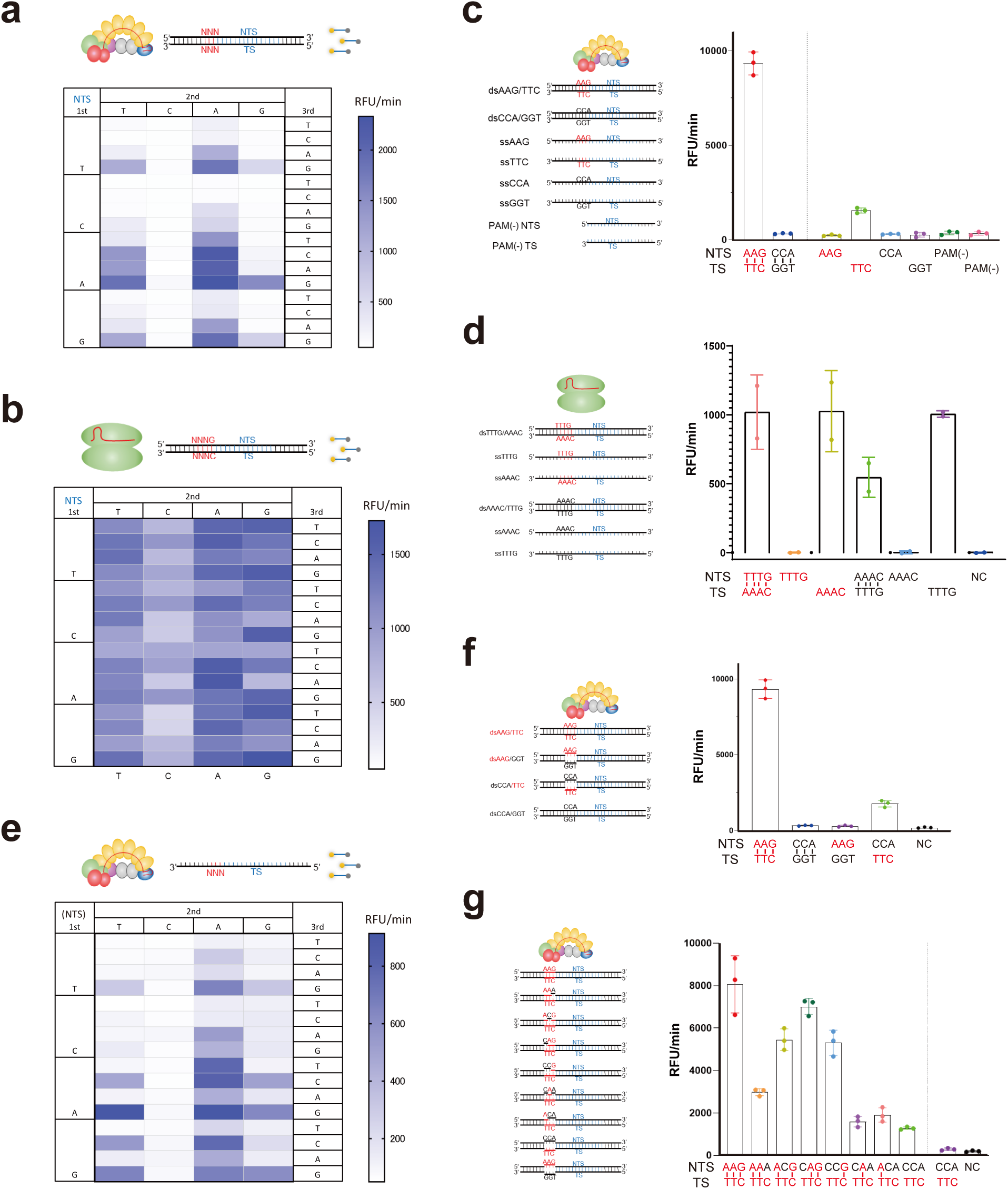
Specificity of PAM recognition for collateral ssDNA cleavage by EcoCas3. (a,b) Screening of all 64 possible target sites containing each of the three-nucleotide PAM sequences for *trans* cleavage activity by EcoCas3 (a) and by LbaCas12a (b). The heat maps represent the RFU per min for collateral cleavage activity. (c) *Trans* ssDNA cleavage by a crRNA-complementary or non-complementary ssDNA (TS or NTS, respectively). EcoCas3/EcoCascade partially activated by TS ssDNA in a PAM-dependent manner (TTC only). (d) LbaCas12a activated by TS ssDNA in a PAM-independent manner (both AAAC and TTTG). (e) Screening of all 64 possible target sites containing each of the three-nucleotide PAM sequences for collateral cleavage activity by the TS ssDNA. (f) Collateral cleavage activated by dsDNA containing an unpaired PAM. (g) Screening of PAM base-pairing between every three PAM nucleotides for *trans* cleavage activity.

According to previous reports ^40, 47, 48^, binding of the ssDNA complementary to the crRNA activates Cas12a for nonspecific *trans* cleavage. We also observed that EcoCas3 and LbaCas12a were activated by crRNA-complementary ssDNA (TS) but not by non-complementary ssDNA (NTS) (Fig. 2c,d). However, the PAM specificity was different between EcoCas3 and LbaCas12a. LbaCas12a was activated by both crRNA-complementary TS flanked by a PAM (AAAC) or a nonPAM (TTTG) (Fig. 2d), as previously reported ^40, 47^. In contrast, EcoCas3 was partially activated by a TS with a PAM (TTC) but a TS with a nonPAM (GGT) prevented any activity (Fig. 2c). We then tested TS PAM specificity for all 64 possible target sites (Fig. 2e). The PAM specificities for ssDNA-activated collateral cleavage were similar to those of dsDNA-activated collateral cleavage (Fig. 2a), although the activity was mostly lower for ssDNA-activated cleavages, except for when the third nucleotide of the PAM was C, such as TAC, AGC, GTC, GAC, and GGC, when the relative fluorescence was increased (Fig. 2e).

Base-pairing between the TS and NTS leads to correct Cascade binding of the NTS, accessibility of the EcoCas3 cleavage site, and degradation of the target DNA ^23, 49^. We observed that dsDNA containing an unpaired PAM between NTS-nonPAM (CCA) and TS-PAM (TTC) partially activated EcoCas3 for collateral cleavage (Fig. 2f). This is not consistent with a previous report that showed dsDNA with an unpaired PAM did not activate EcoCas3 to degrade target DNA substrates ^23^. Screening of PAM base-pairing between each of the three nucleotides showed that base-pairing of the third nucleotide positively affected collateral cleavage activity, and that base-pairing of the first and second nucleotides additively increased the activity of the third nucleotide base-pairing (Fig. 2g). Together, these results of PAM recognition specificity are mostly consistent with results from *in vitro* reconstitution ^23, 49^ and of crystal structure analysis ^18^, except for the partial activity detected for collateral cleavage, in contrast to no activity for target DNA degradation by unpaired PAM recognition ^23^.

### EcoCas3 cleaves the NTS in *cis* followed by the TS in *trans* in a helicase-dependent manner

Complete R-loop formation by the Cascade/crRNA complex recruits the Cas3 helicase/nuclease, which repeatedly cleaves the NTS via the HD domain’s single catalytic site ^31, 32^. It remains unknown how EcoCas3 cleaves the TS and progressively degrades the dsDNA substrate (Fig. 1a). Considering the collateral non-specific ssDNA cleavage in *trans*, we hypothesized that the TS can be cleaved in *trans,* following *cis* cleavage of the NTS after target dsDNA unwinding by the helicase properties of Cas3. To test this, we designed fluorescently-labeled target dsDNA substrates, 5′-NTS-FAM, and 5′-TS-TAMRA, to visualize dsDNA cleavage by EcoCas3 (Extended data Fig. 9a). In control experiments, SpCas9 cleaved both NTS and TS at 3–4 nucleotides upstream of the PAM site, as expected (Fig. 3a). In contrast, the highest peak of EcoCas3 cleavage was 10–11 nucleotides downstream of the PAM site on the NTS, while several peaks upstream of the PAM site demonstrated repetitive cleavage of the NTS. We also observed repetitive cleavage of dozens of nucleotides upstream of the TS PAM, which was likely reeled by EcoCas3 helicase activity and cleaved by its *trans* cleavage activity (Fig. 3a).

**Figure 3.**
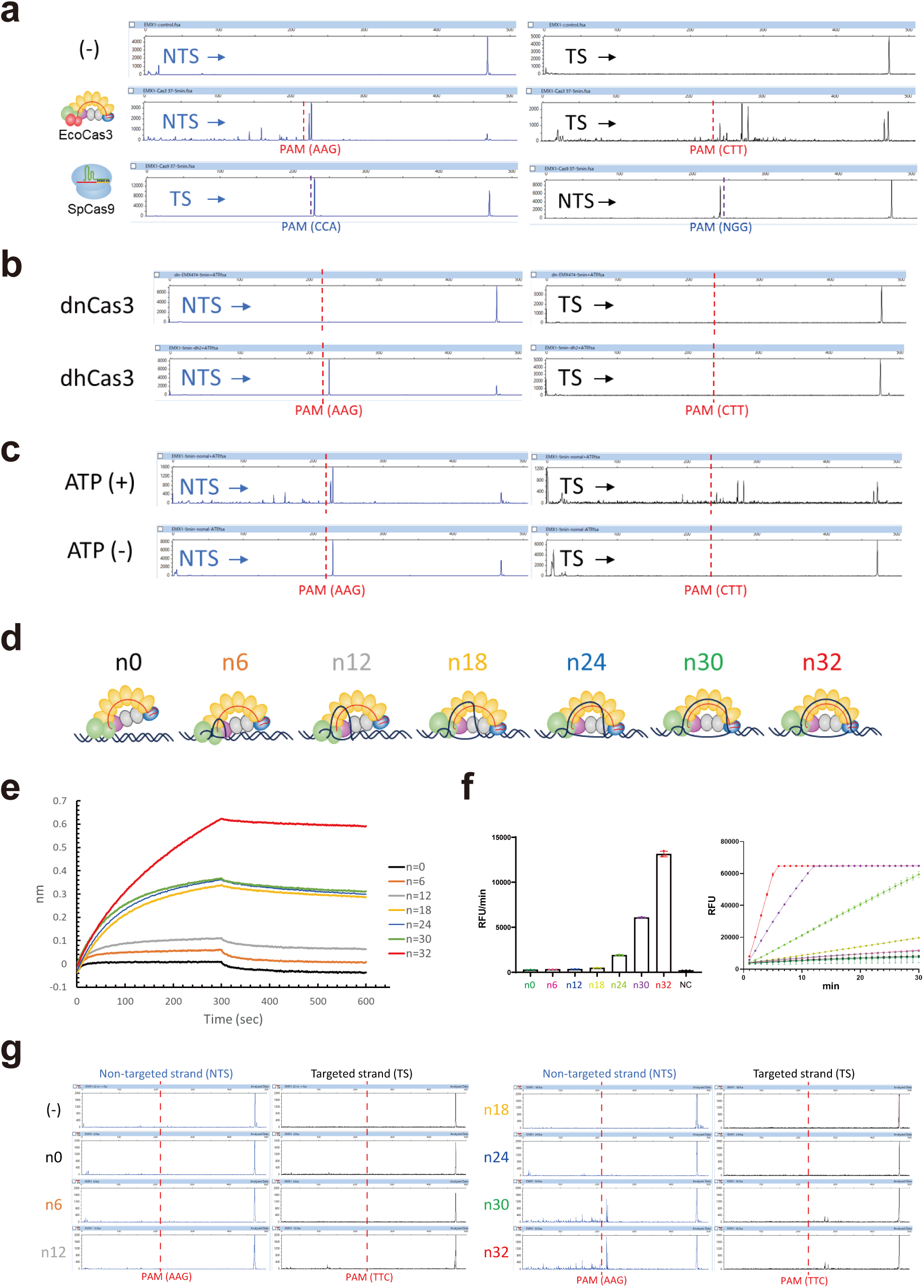
Mechanistic insight into collateral ssDNA cleavage and target DNA degradation. (a) Fluorescently-labeled target dsDNA substrates, 5′-NTS-FAM, and 5′-TS-TAMRA, to visualize dsDNA cleavage. EcoCas3 with EcoCascade RNPs cleaves at NTS nucleotides 10–11, downstream of the PAM site, with repetitive cleavage. The TS cleaved repetitively dozens of nucleotides upstream of the PAM. SpCas9 cleaves both NTS and TS at 3–4 nucleotides from the PAM. (b) The dnCas3 HD domain mutant and the dhCas3 SF2 domain mutant for the dsDNA cleavage assay. (c) The dsDNA cleavage assay by EcoCas3 in ATP (+) or ATP-free (-) reaction buffer. (d) Changing the size of the R-loop formation from 0 to 32 nucleotides by adding multiples of six nucleotides. (e) Measurement of EcoCascade-target DNA associations and dissociations in real-time using a bio-layer interferometry (BLI) biosensor (Octet RED 96 system). (f) In the collateral cleavage assay, n0–n12 base-pair hybridization did not show any cleavage activity, while n18–n32 R-loop formations increasingly promoted *trans* ssDNA cleavage activity. (g) In the dsDNA cleavage assay, the n0–n24 R-loop formation did not produce any cleavage of the NTS or TS. The 30–32 base-pair R-loop formations underwent repetitive cleavage on the NTS and the TS of the target dsDNA substrates.

To confirm the NTS and TS cleavages mediated by nuclease/helicase activities of EcoCas3, we tested a dnCas3 HD domain mutant and a dhCas3 SF2 domain mutant in the dsDNA cleavage assay (Fig. 3b). The dnCas3 mutant cleaved neither NTS nor TS, indicating that the single catalytic domain of EcoCas3 plays a role in generating double-strand breaks (DSBs). Notably, the dhCas3 mutant cleaved the NTS in *cis*, but not the TS in *trans* (Fig. 3b), which was not consistent with the assay’s collateral cleavage results where the dhCas3 mutant cleaved non-specific ssDNA in *trans* (Fig. 1f). The dsDNA cleavage assay for wild-type EcoCas3 and dhCas3 mutant in ATP-free reaction buffer resulted in *cis* cleavage of the NTS, but no *trans* cleavage of the TS (Fig. 3c and Extended data Fig. 9b). Together, these results indicate that the dhCas3 mutant (S483A/T485A) works as an EcoCas3 Nickase and that the helicase activity of EcoCas3 is indispensable not only for repetitive *cis* cleavage of the NTS, but also for *trans* cleavage of the reeled TS.

To further characterize *cis* and *trans* cleavage by EcoCas3, we compared 30 sec (short) and 5 min (long) incubation times for the dsDNA cleavage assay. More prolonged incubation increased repetitive cleavage of the NTS in *cis* and the TS in *trans* (Extended data Fig. 10a). We also observed that progressive *cis* and *trans* cleavages showed similar patterns in the repetitive experiments and the short and long incubation experiments, depending on the target DNA sequence (Fig. 3 and Extended data Fig. 10a). The sizes of many cleaved fragments were between 30–60 bps, which may be used for CRISPR adaptations as previously reported ^12, 50^ (Fig. 3 and Extended data Fig. 10b).

### Incomplete binding of EcoCascade to target DNA with tolerated mismatches elicits collateral ssDNA cleavage but not target dsDNA degradation

We previously reported that a single mismatch within the seed region markedly affected target DNA degradation in the EcoCascade/Cas3 system ^11^. We therefore investigated the effect of mismatch for each nucleotide in the 32-nt spacer on collateral ssDNA cleavage activity. A single mismatch in the spacer region, even within the seed region (positions 1–8), resulted in little or no effect on collateral cleavage activity (Extended data Fig. 11a,b). In the LbaCas12a system, 1–3 mismatches in the seed region also did not affect collateral cleavage activity (Extended data Fig. 11c), consistent with previous reports ^48, 51^. Previous *in vitro* analysis revealed the effect of single mismatches in the target sequence, which slow the rate of R-loop formation and target-strand cleavage by Cas12a ^52, 53^. To investigate whether Cascade-binding and R-loop-formation are linked with collateral cleavage and target DNA degradation, we sought to characterize Cascade-target DNA binding kinetics using a Bio-layer interferometry (BLI) biosensor ^54^. Corresponding to the collateral cleavage assay results (Fig. 2c), crRNA-complementary TS-ssDNA showed associations with EcoCascade but not with non-complementary NTS-ssDNA (Extended data Fig. 12a and Supplementary Table 1). Notably, the crRNA-complementary TS-PAM (TTC) showed higher association than that of TS-nonPAM (GGT) or -PAMless (Extended data Fig. 12a). Moreover, dsDNAs containing a paired PAM (AAG-TTC) showed the maximum EcoCascade-target DNA binding (Extended data Fig. 12b and Supplementary Table 1), which corresponds to the results of the collateral cleavage assay (Fig. 2f). Unpaired PAM between TS-PAM (TTC) and NTS-nonPAM (CCA) indicated a lower association, and unpaired PAM between NTS-PAM (AAG) and TS-nonPAM (GGT) showed little association (Extended data Fig. 12b). Taken together, BLI can provide solid information on the affinity and stability of interactions as previously reported ^54^.

To further investigate the relationship between the R-loop-formation and EcoCas3-mediated collateral ssDNA cleavage and dsDNA degradation, we assayed different length R-loop formations, from 0–32 nucleotides (n0, n6, n12, n18, n24, n30, and n32) (Fig. 3d). BLI revealed that crRNA-DNA hybridization with 0–12 base-pairs (n0, n6, n12) including seed sequences did not show any association, while 18–30 base-pairs (n18, n24, n30) produced a degree of association. Furthermore, a complete match for 32 base-pairs (n32) resulted in stable and emphatic Cascade binding, similar to locked R-loop formation reported previously ^15, 19, 28^ (Fig. 3e and Supplementary Table 1). In the collateral cleavage assay, n0, n6, and n12 did not show any cleavage activity, while n18–n32 R-loop formations increasingly promoted *trans* cleavage of ssDNA (Fig. 3f). In the dsDNA cleavage assay, n0–n24 did not show any cleavage of either NTS or TS DNA (Fig. 3g). This means that collateral cleavage does not need the nicking activity on the NTS (n18 and n24). Only the n30 and n32 sequences underwent repetitive cleavage on both the NTS and TS, and progressive cleavage of target dsDNA substrates (Fig. 3g). Taken together, these results show two Cascade binding modes. Intermediate R-loop formation by mismatches on the spacer sequences elicits collateral ssDNA cleavage. Complete R-loop formation with full crRNA-DNA hybridization leads to repetitive *cis* cleavage of the NTS with *trans* cleavage of the TS to degrade the target dsDNA substrate, as described in previous reports ^15, 19, 28^.

### Dynamic visualization of CRISPR interference: PAM search, nicking, and DSB

Cryo-EM and smFRET are not capable of visualizing how EcoCas3 degrades target dsDNA ^12, 30^. We therefore employed hs-AFM, which enables real-space and real-time observations at the macromolecule level, as previously shown by visualizing CRISPR-Cas9 interference ^55^. First, we visualized the binding of Cascade/crRNA to a target DNA, a 645-bp dsDNA containing a target spacer site flanked by a PAM (AAG) at 219-bp and 423-bp from the ends of the DNA fragment (Fig. 4a). We then adsorbed the mixture of donor DNAs and EcoCascade RNPs onto a 3-aminopropyl-trietoxy silane-mica surface (APTES-mica) ^56^. We observed that the EcoCascade RNP ran from one end to the other through the target DNA, presumably searching for the right PAM site and spacer sequences (Fig. 4b and video 1 and 2). We also found that many EcoCascade RNPs formed a stable multibody and stuck to the expected target site. Notably, we observed a typical DNA bend at the EcoCascade-RNP-binding site for stable R-loop formation, as previously indicated by cryo-EM ^15, 18^ and smFRET studies ^16, 57^. During the observation periods, the EcoCascade RNPs bound tightly to the target DNAs without dissociating, consistent with previous smFRET analyses ^16, 20, 57^. By applying excessive force, the EcoCascade RNP body was broken and separated into multiple Cas effector components (Extended data Fig. 13a and video 3).

**Figure 4.**
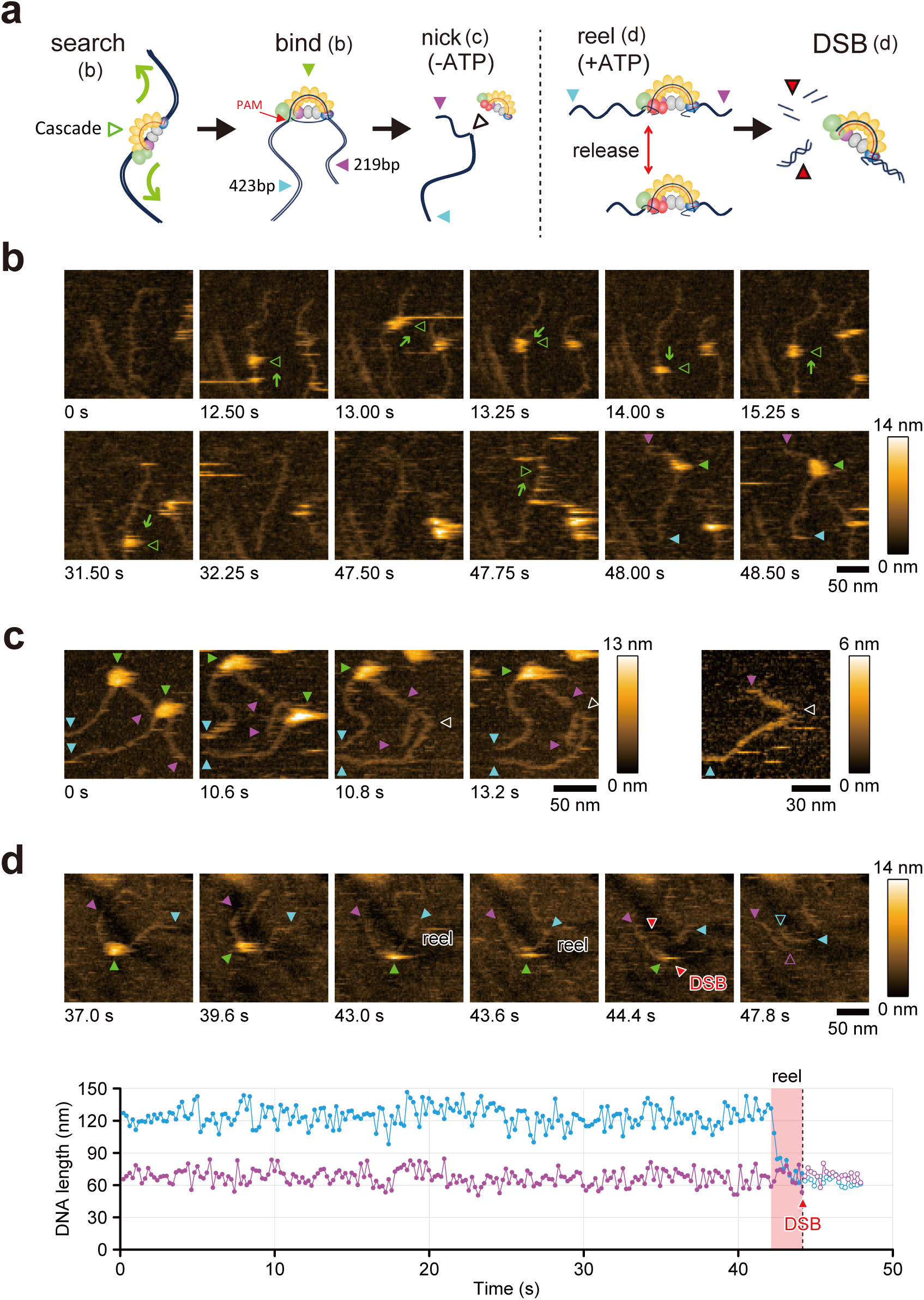
Dynamic visualization of type I CRISPR interference by hs-AFM. (a) Schematic depictions of type I CRISPR interference (b–d). (b) High-speed atomic force microscopy (hs-AFM) visualizes the EcoCascade RNP (green triangle) searching for an appropriate PAM site from one end of the target DNA to the other (green arrows) (video 1). EcoCascade binds to the target DNA, a 674-bp dsDNA containing a target spacer site flanked by PAMs (AAG) at 219-bp and 423-bp from the ends of the DNA fragment (purple and blue triangle, respectively). (c) Injection of EcoCas3 protein after fixing EcoCascade RNPs with the target DNA in ATP-free (-) reaction buffer to produce EcoCas3-mediated nicking (white triangle) at the target site (video 4). (d) In the ATP (+) reaction buffer, the EcoCas3-EcoCascade complex reels the longer side of the DNA (blue triangle) and then cleaves it with a DSB (red triangles) (videos 6).

Next, we injected EcoCas3 proteins after fixing the EcoCascade RNPs with the 645-bp target DNA in ATP-free reaction buffer to reproduce EcoCas3-mediated nicking at the target site. EcoCas3 did not make any single-strand breaks (SSBs) *per se* but together with the Cascade RNPs, several SSB-like DNA bends at the target site were observed (Fig. 4c). Notably, the shape of DNA bending was similar to that of artificially nicked dsDNA using Nb.BsrDI nicking endonucleases (Extended data Fig. 13b). Furthermore, we observed dynamic movement of the EcoCas3-Cascade RNP along the target DNA, which suddenly bound to the target site and disconnected from the DNA, with the bent DNA appearing as an SSB shape (Fig. 4c and video 4). In contrast, in ATP-containing reaction buffer, we detected many DNA fragments of 219-bp or 423-bp after injection of EcoCas3 proteins. Notably, we observed the EcoCas3-Cascade complex bound to the target site was repeatedly reeling the longer side of the DNA then releasing it (Extended data Fig. 14 and video 5). Finally, we captured the dynamic movements by which the EcoCas3-Cascade complex shortened the target DNA and cleaved it with a DSB, followed by release of the DNA from the EcoCas3-Cascade complex (Fig. 4d and video 6 and 7).

## Discussion

Up until now it has been unclear how a single HD nuclease domain in Cas3 can cause DSBs at target sites and long-range unidirectional deletions upstream of target sites ^10, 11, 13, 15, 25, 29, 30^. We believe that this is the first report to use hs-AFM to capture the dynamic movements of CRISPR-Cas3 interference at the single molecule level. The hs-AFM results clarify that the EcoCascade/crRNA complex searches for and binds to target DNA, and recruited EcoCas3 bound to EcoCascade then reels and loops the target dsDNA, and subsequently cleaves it (Fig. 5). This is consistent with a reeling model in which Cas3 remains associated with Cascade to cleave ssDNA by a reeling mechanism ^12^. However, it remained unknown how EcoCas3 cleaves the reeled TS and progressively degrades the dsDNA substrate (Fig. 1a). Our results from collateral ssDNA cleavage assays and dsDNA cleavage assays revealed that Cas3 repeatedly cleaves the NTS by helicase activity in *cis*. Simultaneously, the TS reeled by the helicase property of Cas3 can be cleaved by non-specific ssDNA cleavage activity in *trans*. The hs-AFM analysis also revealed that Cascade-bound Cas3 repeatedly reels and releases the target DNA upstream of the PAM site, followed by target degradation. (Extended data Fig. 14). Although these results are inconsistent with a translocation model (Fig. 1a), Cas3 with Cas1 and Cas2 forming a primed acquisition complex may translocate in search of protospacers, which was revealed by smFRET ^20, 58^. Despite hs-AFM being able to analyse regions of hundreds of bp, it is not feasible to visualize long-range dsDNA cleavages (in the order of kb) *in vitro*. Previous *in vivo* experiments with the CRISPR-Cas3 system ^10, 11^ indicated unidirectional long-range deletions, where the spacer sequences and the PAM site remained in the absence of indel mutations and repetitive fragmented DNA deletions upstream of the PAM sites. The hs-AFM movement capture results support this CRISPR interference phenomenon, where the EcoCascade/Cas3 complex remains bound to the target site to repeat DNA degradation, which may expand the deletion size.

**Figure 5.**
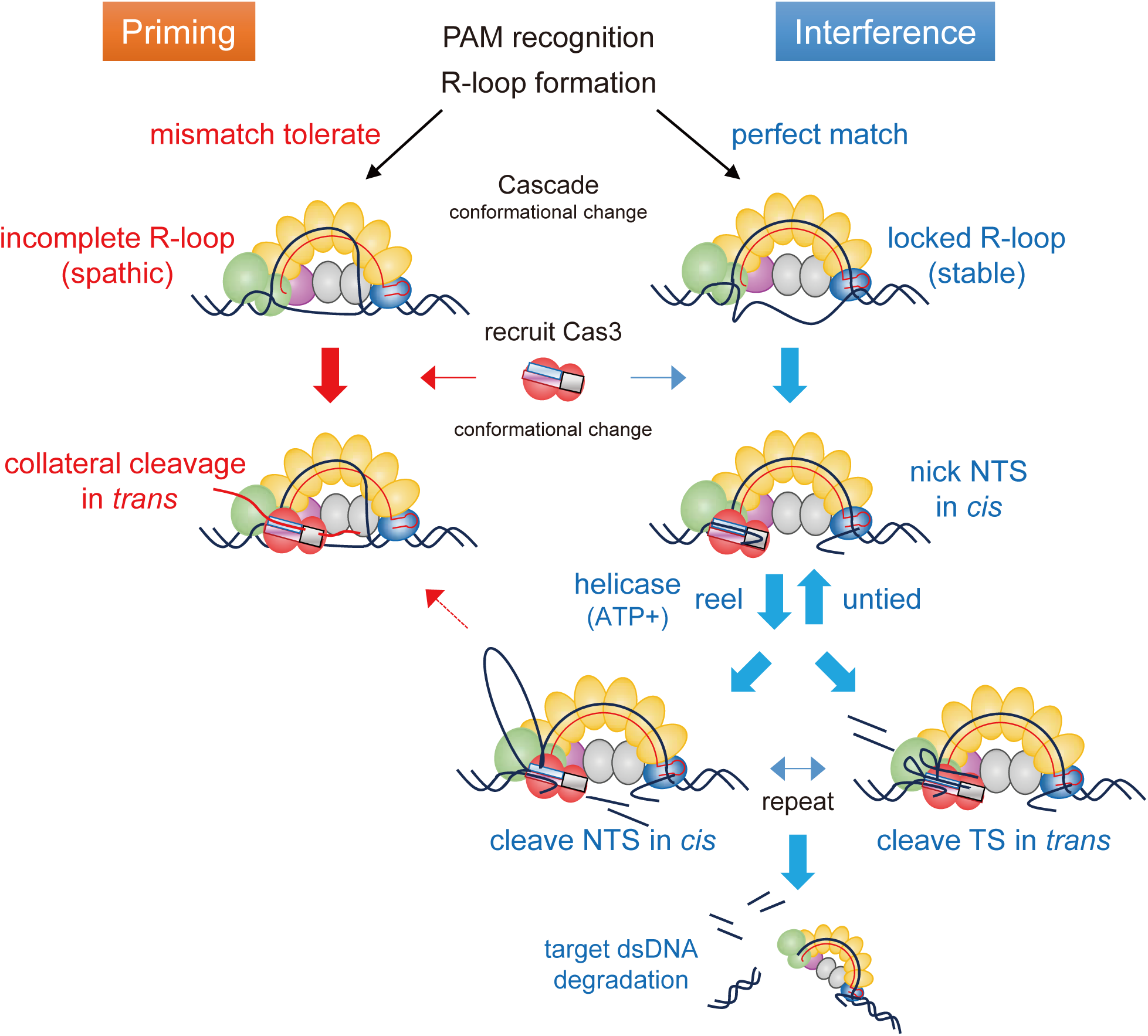
Mechanism of type I CRISPR interference and priming. Cascade binding to target DNA with tolerated mismatches elicits collateral ssDNA cleavage as a priming mode. Stable Cascade binding via a complete R-loop formation drives nicking of the non-target strand (NTS), followed by helicase-dependent unwinding of double-stranded DNA (dsDNA) upstream of the PAM site. Then, *trans* cleavage of the target strand (TS) combined with repetitive *cis* cleavage of the NTS degrades the target dsDNA substrate as an interference mode.

Previous smFRET studies revealed that type I CRISPR systems have two binding modes for target recognition called interference and priming ^19, 20, 57, 59^ (Fig. 5). The low fidelity priming mode allows a whole range of mutated invaders to be detected and the priming process to be initiated ^19, 48, 51, 57^. Meanwhile, the high-fidelity interference mode ensures the destruction of perfectly matching targets to destroy foreign invaders without new spacer acquisition. Our results using the BLI biosensor show that partial R-loop formation (18–24 bp) enforces short-lived and unstable Cascade binding, whereas full R-loop formation (30 and 32 bp) provides locked and immobilized Cascade binding to the target DNA (Fig. 3e). We also find that this partial Cascade binding can recruit Cas3 to mediate non-specific ssDNA cleavage in *trans*, but can also interrupt dsDNA cleavage in *cis* (Fig. 3f,g). Moreover, this collateral ssDNA cleavage tolerates mismatches within the spacer sequences (Extended data Fig. 11a,b), in contrast to our previous findings with target dsDNA cleavage ^11^. Previous *in vivo* studies have revealed that the CRISPR-Cas system acquires new spacer sequences from escape mutants that carry mutations in PAM and protospacer sequences, known as primed CRISPR adaptation or priming ^19, 20, 57, 58^. These findings suggest that the type I CRISPR system uses the collateral ssDNA cleavage for the priming process (Fig. 5). This is also supported by recently reported results showing that Cas12a has multiple nicking activities with tolerance of 4–8 mismatches within the PAM and spacer sequences in a natural role as an immune effector against rapidly evolving phages ^48, 51^.

Our results also indicate that only stable Cascade binding can initiate nicking of the NTS and activate the ATP-dependent helicase property of Cas3, which reels and loops the TS for cleavage in the interference mode ^15, 19, 28^ (Fig. 5). It is still unclear what the critical step is that drives the Type I CRISPR interference mode. Previous cryo-EM studies reveal that full R-loop formation following conformational changes of Cascade triggers a flexible bulge in the NTS, enabling Cas3 nicking in this region ^15, 30^. In addition, the *trans* cleavage activities can be controlled by multiple steps including specific PAM recognition, R-loop formation-dependent conformational changes in Cascade, and recruitment of Cas3 and conformational changes in Cas3 itself, as previously suggested by several cryo-EM studies ^15, 17, 18, 21, 22, 26, 60^. A chip-hybridized-association-mapping-platform (CHAMP) that measures interactions between proteins and DNA sequences also indicated that Cas3 recruitment is sensitive to PAM and PAM-proximal DNA-RNA-mismatches ^61^. It also showed that DNA sequence-specific loss of Cas8 abrogates Cas3 recruitment and provides an additional proofreading mechanism for modulating CRISPR interference ^61^. These proofreading mechanisms of type I CRISPR are similar to those of other proofreading systems, such as the conformational checkpoint of the HNH nuclease domain in type II Cas9 ^62, 63^ and the potential lid movement of the RuvC catalytic domain in type V Cas12 ^64, 65^.

Crystal structure analysis of TfuCas3 ^30, 66^ showed that EcoCas3 recruited to Cascade may also be activated by guiding ssDNA to the HD domain by either of two routes, the bypass route or the helicase tunnel. Considering our findings and crystal structure data ^30, 66^, EcoCas3 may continuously cleave NTS via the helicase tunnel ^30^. Simultaneously, EcoCas3 cleaves the TS by collateral *trans* cleavage activity through the bypass route. By repeating *cis* NTS cleavage via the helicase tunnel route and *trans* TS cleavage via the bypass route, EcoCas3 may achieve phage plasmid degradation in *E. coli* ^8^ and large-scale genome editing in human cells ^10, 11^ (Fig. 5).

The CRISPR-Cas3 system potently degrades phage and viral DNA. It is probably more powerful than Cas9 and Cas12, which carry small mutations ^2–4^. However, if Cas3 is too powerful, it may have the potential for self-attack, from which Cas3 must escape. EcoCas3 has a longer spacer sequence of 27 nucleotides compared with the 20 nucleotides of Cas9 or the 24 nucleotides of Cas12, which may increase the specificity for target recognition. EcoCas3 has maximal cleavage activity at 37°C, although EcoCas3 protein is sensitive to temperature-dependent aggregation at 37°C (Extended data Fig. 3), which may also decrease self-attack. The specific PAM recognition by EcoCascade can also enable escape from self-attack (Fig. 2) because various CRISPR-Cas systems have a PAM system that distinguishes self from non-self ^27^. Crystal structure analysis showed that Cas9 and Cas12 enable R-loop formation by first recognizing and unwinding the NTS-PAM ^27^. Compared with Cas9, Cas12a loosely fits the PAM binding channel, allowing it to slightly open during suboptimal PAM binding ^67^. The resulting loss of specific interactions between the PAM and the Cas12a channel can explain the observed higher *trans* cleavage activities of Cas12a ^67^. In the CRISPR-Cas3 system, EcoCas3 recruitment and binding to EcoCas8 depend on TS-PAM recognition. EcoCas8 binds to the third position of the TS-PAM and unwinds through recognition of the NTS-PAM ^18^, which may increase PAM specificity in collateral ssDNA cleavage (Fig. 2). In the CRISPR-Cas3 system, the PAM plays an important role in self- and non-self-discrimination, and PAM recognition by Cas effectors is the initial step following the formation of an R-loop structure with the crRNA ^18, 27^.

In conclusion, we found that the partial binding of EcoCascade to target DNA can elicit collateral non-specific ssDNA cleavage in the priming mode. With stable binding by a complete R-loop formation, the collateral ssDNA cleavage can be used for *trans* cleavage of the TS combined with repetitive *cis* cleavage of the NTS to degrade the target dsDNA substrate in the interference mode. These results provide a mechanistic insight into collateral ssDNA cleavage and target DNA cleavage by the CRISPR-Cas3 system, enabling further understanding of type I CRISPR priming and interference of foreign DNAs.

## Acknowledgments

We thank T. Omoto, and S. Kobori at Osaka University, M. Hoshi, K. Mochizuki, and A. Fukui at Tokyo University, and S. Saji, S. Yamamoto and N, Godai at the RIKEN SPring-8 Center for their technical assistance and Drs. T. Ando and T. Watanabe-Nakayama for technical support with HS-AFM. We also thank Professor Jacob Corn at ETH Zurich for scientific advice and Jeremy Allen, PhD from Edanz Group (https://en-author-services.edanz.com/ac) for editing a draft of this manuscript. This project was supported by JSPS KAKENHI Grant Number 18H03974 (T.M.), 19K16025 (K.Y.), 19KK0401 (K.Y.) and 20H00327 (N.K.) and JST-CREST (JPMJCR1762 to N.K.). Support also came from the Platform Project for Supporting Drug Discovery and Life Science Research [Basis for Supporting Innovative Drug Discovery and Life Science Research (BINDS)] from AMED under Grant Number JP20am0101070 (support numbers 1251 and 2463).

## Author Contributions

K.Y. and S.S. designed and performed most of the experiments, analyzed the data with assistance from Y.K. and Y.Y. K.T., M. O. and M.Y. prepared and characterized the CRISPR-Cas proteins. N.K., K.Y. and K.T. performed the hs-AFM experiments. T.M. conceived and supervised the study, and wrote the manuscript with editorial contributions from all authors.

## Competing interests

S.S. and Y.K. are employees of C4U. K.Y., K.T. and T.M. are scientific advisors for C4U. The other authors declare no competing interests.

## Data availability

All data supporting the findings of this study are available from the corresponding author on reasonable request. Source data are provided with this paper.

## Methods

### Expression and purification of EcoCas3 and EcoCascade/crRNA

We employed a method to express recombinant EcoCas3 at a low temperature using a baculovirus expression system. Briefly, we cloned an EcoCas3 cDNA with a octa-histidine tag and a six asparagine-histidine repeat tag into a pFastbac-1 plasmid (Thermo Fisher Scientific, Waltham, Massachusetts, USA) according to the manufacturer’s instructions (Extended data Fig. 2a). The TEV protease recognition site was also inserted between the tags and EcoCas3 to enable tag removal. Self-ligation of the PCR product generated the mutant Cas3, such as H74A (dead nickase; dn) and S483A and T485A double mutant (dead helicase; dh) with PrimeSTAR MAX (Takara Bio, Kyoto, Japan). Coding sequences cloned in the plasmids are listed in Supplementary Table 2.

Expression of EcoCas3-tag fusion proteins in Sf9 cells. We infected Sf9 cells with baculovirus at a multiplicity of infection (MOI) of two at 28°C for 24 h. Then, we changed the culture temperature to 20°C four days after infection for protein expression. Sf9 cells were then collected and stored at -80°C until use. The expressed EcoCas3-tag fusion proteins were purified using nickel affinity resin (Ni-NTA, Qiagen, Hilden, Düsseldorf, Germany). To remove tags, purified protein was digested with TEV protease and then further purified by size-exclusion chromatography using Superdex 200 Increase 10/300 GL (Thermo Fisher Scientific) in 0.2 M NaCl, 10% glycerol, 1 mM DTT, and 20 mM HEPES-Na (pH 7.0).

Cascade from *E. coli* and CRISPR RNA complex (EcoCascade/crRNA) was produced as described previously ^23, 38^. Briefly, we cloned EcoCas11 with a hexahistidine tag and HRV3C protease recognition site, EcoCascade operon, and pre-crRNA into pCDFDuet-1, pRSFDuet-1, and pACYCDuet-1 plasmids, respectively (Extended data Fig. 2c). Sequences cloned in these plasmids are also listed in Supplementary Table 2. Then, we transformed JM109(DE3) with three plasmids to express EcoCascade/crRNA recombinant protein complex. Expressed recombinant EcoCascade-crRNA was purified using Ni-NTA resin. After removal of the hexahistidine tag by HRV3C protease, EcoCascade-crRNA was further purified by size-exclusion chromatography in 350 mM NaCl, 1 mM DTT, and 20 mM HEPES-Na (pH 7.0).

### Thermal stability assay of EcoCas3

Thermal stability was evaluated by nanoDSF using the Tycho NT.6 system (NanoTemper Technologies GmbH, München, Germany) ^68^. Also, Thermal stability at a constant 37°C was measured by a thermal shift assay using a Mx3000p real-time PCR instrument (Agilent technologies, Santa Clara, California, USA) and SYPRO orange (Thermo Fisher Scientific) ^69^.

### Single and double-stranded DNA preparation

To detect *in vitro* DNA cleavage activity of CRISPR-Cas3 proteins, targeted sequences of *EMX1* with PAM variants (AAG or CCA) were cloned into a pCR4Blunt-TOPO plasmid vector (Thermo Fisher Scientific) according to the manufacturer’s protocol. For collateral DNA cleavage assays, 60 bp activator fragments of *hEMX1* and *mTyr* (which included a target site) were designed and purchased. Targeted sequences for CRISPR-Cas3, CRISPR-Cas12a and CRISPR-Cas9 are listed in Supplementary Table 3. PAM sequence variants and targeted sequence variants were also designed to examine collateral ssDNA cleavage activity. Biotin-labeled fragments were also purchased for protein-DNA interaction analysis. For fragment analysis, fluorescence-labeled primers were designed and the DNA fragment amplified from genomic DNA of HEK293T cells using Gflex DNA polymerase (Takara-bio). Amplicons were purified using NucleoSpin Gel and a PCR Clean-up kit (Takara-bio) according to the manufacturer’s protocols. A DNA fragment for hs-AFM was also amplified with non-labeled primers. All sequences of primers and donor DNAs are listed in Supplementary Table 4 and 5, respectively.

### *In vitro* DNA cleavage activity

To analyze DNA cleavage activity, 1.6 nM of plasmid with or without targeted sequences were added to 115 nM EcoCascade-crRNA complex, 250 nM EcoCas3, and 2.5 mM ATP in CRISPR-Cas3 working buffer (60 mM KCl, 10 mM MgCl_2_, 10 µM CoCl_2_, 5 mM HEPES-KOH, pH 7.5), as previously described ^40, 41, 45^. After incubation at 37°C, samples were detected by either electrophoresis or with the MultiNa microchip electrophoresis system and the DNA-12,000 kit (Shimadzu, Kyoto, Japan).

### Reporter assay for DNA and RNA cleavage

To characterize Cas3 collateral nucleic acid cleavage activities, 50 nM DNA activator templates were added to 100 nM EcoCascade-crRNA complex, 250 nM EcoCas3 and 2.5 mM ATP in CRISPR-Cas3 working buffer (60 mM KCl, 10 mM MgCl_2_, 10 µM CoCl_2_, 5 mM HEPES-KOH, pH 7.5). We used the DNase Alert kit (Integrated DNA Technologies, Coralville, IA USA) and the RNase Alert kit (Integrated DNA Technologies) for detecting DNase and RNase activity, respectively. To measure the ssDNA cleavage activity, we used the qPCR reporter probe for GAPDH (the sequence is listed in Supplementary Table 4) at 125 nM. Cleavage-related change in fluorescence signal of the probe was measured every 30 s for 60 min under incubation at 37°C using a Real-time PCR system (Bio-Rad Laboratories, Hercules, California, USA). Alternatively, M13mp18 single-stranded DNA (New England Biolabs, Ipswich, Massachusetts, USA) or pBluescript plasmid were added and incubated at 37°C. Samples were then electrophoresed on an agarose gel.

### DNA fragment analysis

To analyze CRISPR DNA cleavage patterns *in vitro*, 16 nM DNA fragments amplified from HEK293 genomic DNA were added to 160 nM EcoCascade-crRNA complex, 400 nM EcoCas3 and 2 mM ATP in CRISPR-Cas3 working buffer (60 mM KCl, 10 mM MgCl_2_, 10 µM CoCl_2_, 5 mM HEPES-KOH pH 7.5). After incubation at 37°C, DNA samples were purified by ethanol precipitation. The length of DNA in samples was measured using GeneScan 600 LIZ dye Size Standard (Thermo Fisher Scientific) via a G5 dye set filter. All data were analyzed using PeakScanner software (Thermo Fisher Scientific).

### Protein-DNA interaction assay

The evaluation of binding properties between EMX-EcoCascade (analyte) and target DNAs (ligands) was performed by bio-layer interferometry (BLI) using the Octet RED 96 system (ForteBio, Sartorius BioAnalytical Instruments, Fremont, California, USA). All ligands were biotinylated (20 µM) and immobilized on streptavidin biosensors. Kinetic titration series were performed in interaction buffer (PBS with 0.01% Tween 20, 0.02% BSA). Analyte concentration was 20 µM in the interaction buffer. The association and dissociation times were both 300 sec to measure the interaction between ligands and analyte. These raw data were analyzed using ForteBio analysis software. The binding sensorgram was locally fitted to a 1:1 Langmuir binding model with mass transport limitation. Sequences for the donor DNA fragments were listed in Supplementary Table 5.

### High-speed atomic force microscopy (hs-AFM)

hs-AFM imaging was performed in solution using a laboratory-built hs-AFM setup as described previously ^55^. We used small cantilevers (BLAC10DS-A2, Olympus, Tokyo, Japan) with a nominal spring constant of 0.1 N/m, resonance frequency of ∼0.5 MHz, and a quality factor of ∼1.5 in the buffer. The cantilever’s free oscillation amplitude *A*_0_ and set-point amplitude were set at 1−2 nm and ∼0.9 × *A*_0_, respectively. To observe either the pre-mixed complex of EcoCascade/crRNA and dsDNA or the artificially nicked dsDNA at high spatial resolution, hs-AFM imaging was carried out in observation buffer (5 mM HEPES-KOH, pH 7.5, 30 mM KCl, 1 mM MgCl_2_, 2 μM CoCl_2_, 10% glycerol) at room temperature (∼25°C) using 3-aminopropyltriethoxysilane treated mica as described previously ^55^.

To observe dynamic behaviors of EcoCascade and EcoCas3 on dsDNA, we used a mica-supported lipid bilayer (mica-SLB) as a sample substrate. To observe EcoCascade binding to a target site, a lipid composition of 90:5:5 (w/w) DPPC:DPTAP:biotin-cap-DPPE was used. We deposited 5 nM dsDNA amplicon on the mica-SLB. Three minutes later the sample surface was rinsed with 20 μl observation buffer. We then immersed the sample stage in a liquid cell containing about 60 μl observation buffer, and hs-AFM imaging was performed in a room heated to ∼30°C with a heater. We added a drop (∼6 μl) of EcoCascade to the liquid cell during the hs-AFM observations, resulting in a final concentration of ∼20 nM. To observe DNA reeling and double-strand break generation by EcoCas3, the lipid composition used was 80:10:10 (w/w) DPPC:DPTAP:biotin-cap-DPPE. EcoCascade-DNA pre-assembled with 20 nM EcoCascade in observation buffer was placed on the mica-SLB together with 2 nM DNA at 37°C for 5 min. The sample surface was rinsed with 20 μl observation buffer and then imaged with hs-AFM, with a head temperature controlled at ∼37°C using a thermostatic cover. During the hs-AFM observations, a drop (∼6 μl) of EcoCas3 and ATP mixture was added to the liquid cell, at a final concentration of ∼100 nM and ∼2 mM, respectively. Primers for the donor DNA amplicons are listed in Supplementary Table 4.

## Extended data Figure Legends

**Extended data Fig. 1. Purification of EcoCas3 and TfuCas3 recombinant proteins by the *E. coli* bacterial expression system.** (a) A limited amount of highly aggregated *E. coli* Cas3 (EcoCas3) protein was purified by size-exclusion chromatography and was evaluated by sodium dodecyl sulfate-polyacrylamide gel electrophoresis (SDS-PAGE). Co-expression of HtpG chaperon and low temperature 20°C culture were used in the *E. coli* expression system. (b) Isolation of large amounts of *Thermobifida fusca* Cas3 (TfuCas3) protein at 37°C.

**Extended data Fig. 2. Purification of EcoCas3 and EcoCascade (Cas5, Cas6, Cas7, Cas8, and Cas11) ribonucleoproteins.** (a) A large amount of EcoCas3 protein was purified using a baculovirus expression system in Sf9 insect cells cultured at 28°C. (b) Purified EcoCas3 protein from Sf9 cells was soluble and ∼95% homogeneous on SDS-PAGE. (c,d) A complex of Cas5, Cas6, Cas7, Cas8, and Cas11 proteins and crRNA was co-expressed in JM109(DE3) *E. coli* cultured at 37°C, purified using Ni-NTA resin, and separated by size exclusion chromatography.

**Extended data Fig. 3. Temperature-dependent stability of recombinant EcoCas3 protein.** To evaluate the temperature-dependent stability of purified EcoCas3, we employed a modified nanoscale differential scanning fluorimetry method, nanoDSF, which determines protein stability by measuring intrinsic tryptophan or tyrosine fluorescence using the Tycho NT.6 system (NanoTemper Technologies GmbH) ^68^. The profiles show the first derivative ratio of 330 and 350 nm as the temperature rises from 35°C to 95°C. EcoCas3 protein purified from Sf9 cells had a temperature inflection point (Ti) of 47.8°C, while TfuCas3 and off-the-shelf SpCas9 proteins had Ti’s of 70.4°C and 49.7°C, respectively. LbaCas12a protein had two Tis (44.8 and 51.9°C), which may represent dimeric structural dissociation between the REC and NUC lobes ^67, 70^.

**Extended data Fig. 4. Temperature-dependent stability of recombinant EcoCas3 protein.** We evaluated temperature-dependent stability using ProteoStat detection reagent (Enzo Life Sciences), which allowed the aggregation onset temperature to be determined ^69^. Vertical and horizontal axes show fluorescence intensity, which is dependent on protein denaturation, and time, respectively. Considering thermal stability at a constant temperature of 37°C, EcoCas3 was mostly denatured in 8 h, while SpCas9, LbaCas12, and TfuCas3 were not denatured after 24 h (upper). At a constant temperature of 48°C, LbaCas12a and SpCas9 proteins showed earlier aggregation onset than EcoCas3 (lower).

**Extended data Fig. 5. Evaluation of R-loop formation by electrophoretic mobility shift assays (EMSAs).** (a) EcoCascade/crRNA complex binds to supercoiled (SC) plasmids containing *hEMX1* spacer sequences flanked by a PAM (AAG) as shown by a red arrow, but did not bind to a nonPAM (CCA). (b) The EcoCascade complex binding to the PCR products of other target sequences (*mTyr* and *rIl2rg* genes). Denaturation of the EcoCascade complex (0.08% SDS, 95°C, 2 min) abolished the DNA binding (black arrows). (c) Exchange of nucleotide pairs between crRNAs and target sequences abolished the binding, indicating the specificity of target recognition by the EcoCascade/crRNA complex.

**Extended data Fig. 6. Collateral cleavage of nearby non-specific ssDNAs**. (a) EcoCas3 possesses collateral non-specific single-stranded DNA (ssDNA) cleavage activity after target-specific double-stranded DNA (dsDNA) cleavage. Circular ssDNA M13 phage and linearized long ssDNA were degraded after incubation for 1 h at 37°C (red arrows), but circular pBlueScript dsDNA was not cleaved (black arrows). (b) Comparison of collateral cleavage activity between 37°C and 20°C. EcoCas3 showed higher collateral cleavage activity at 37°C than at 20°C. (c) *In vitro* reconstitution assay for target DNA degradation also showed higher activity at 37°C (red arrows) than at 20°C (black arrows).

**Extended data Fig. 7. Collateral cleavage activity for ssDNAs and ssRNAs**. (a) Using fluorescent reporter DNA oligonucleotides (DNaseAlert™, IDT), we detected collateral ssDNA cleavage activity by assembling EcoCas3, EcoCascade RNPs and dsDNA fragments that included target sequences (*hEMX1* or *mTyr*) flanked by PAM-AAG or -ATG, but not with PAM-CCA. (b) Little or no RNase activity was detected with fluorescent reporter RNA oligonucleotides (RNaseAlert™, IDT) by assembling EcoCas3, EcoCascade RNPs and dsDNA fragments.

**Extended data Fig. 8. Specific PAM recognition for collateral ssDNA cleavage by EcoCas3.** (a,b) Screening of all 64 possible target sites containing each of the three-nucleotide PAM sequences for *trans* cleavage activity by EcoCas3 (a) and LbaCas12a (b).

**Extended data Fig. 9. The dsDNA cleavage assay.** (a) Schematic depictions of the dsDNA cleavage assay. Fluorescently-labeled target dsDNA substrates, 5′-NTS-FAM and 5′-TS-TAMRA, to visualize dsDNA cleavage. (b) The dhCas3 SF2 domain mutant cleaves the NTS in *cis*, but not the TS in *trans* in ATP (+ and -) reaction buffer, while LbaCas12a cleaves both the NTS and TS in ATP (+ and -) reaction buffer.

**Extended data Fig. 10. The dsDNA cleavage assay.** (a) Comparison of 30 sec (short) and 5 min (long) incubation for the dsDNA cleavage assay (*hEMX1* or *mTyr*). (b) The size and patterns of cleaved fragments after short and long incubation in the dsDNA cleavage assay.

**Extended data Fig. 11. The effect of mismatch of each 32-nt spacer sequence on collateral cleavage activity.** (a,b) A single mismatch in the spacer region has a little or no effect on collateral cleavage activity of *hEMX1* target (a) and *mTyr* target (b). (c) One to three mismatches in the PAM sites and the spacer region have little effect on collateral cleavage activity by LbaCas12a.

**Extended data Fig. 12. EcoCascade-target DNA associations and dissociations measured by the Octet RED 96 System.** (a) crRNA-complementary ssDNA with TS-PAM (TTC) showing collateral cleavage (Fig. 2c) represents higher association than TS-nonPAM or TS-PAMless. (b) The interactions between Cascade and dsDNAs containing paired PAM (AAG-TTC), unpaired PAM between TS-PAM (TTC) and NTS-nonPAM (CCA), and unpaired PAM between NTS-PAM (AAG) and TS-nonPAM (GGT), correspond with the results from the collateral cleavage assay (Fig. 2f).

**Extended data Fig. 13. Dynamic visualization of CRISPR interference by hs-AFM.** (a) By applying excessive force, the EcoCascade RNP body (green triangle) was broken and separated into multiple Cas effector components (white triangle) (video 3). (b) The shape of bent DNA (white arrows) resulting from artificial nicking by Nb.BsrDI endonuclease.

**Extended data Fig. 14. Dynamic visualization of CRISPR interference by hs-AFM.** In ATP (+) reaction buffer, the EcoCas3-Cascade complex repeatedly reels and releases the longer side of the DNA (blue arrows) and then cleaves it with a DSB (red arrows) (video 5).

**Video 1.** EcoCascade RNP searches for the right PAM site from one end to the other through the target DNA (Fig. 4b).

**Video 2.** EcoCascade RNP sticks to the target site of the dsDNA to form R-loop architecture.

**Video 3.** Excessive forces disrupt EcoCascade RNP binding and separate the body into multiple Cas effector components (Extended data Fig. 13a).

**Video 4.** EcoCas3 combined with EcoCascade RNPs mediates nicking at the target site of the 600-bp DNA in ATP-free (-) reaction buffer (Fig. 4c).

**Video 5.** EcoCas3-Cascade complex repeatedly reels and releases the longer side of the DNA, then cleaves it with a DSB in ATP (+) reaction buffer (Extended data Fig. 14).

**Video 6.** EcoCas3-Cascade complex reels the longer side of the DNA, then cleaves it with a DSB in ATP (+) reaction buffer (Fig. 4d).

**Video 7.** EcoCas3-Cascade complex reels the longer side of the DNA, then cleaves it with a DSB in ATP (+) reaction buffer.

