## Supplementary Figures for "Dynamic mechanisms of CRISPR interference by *Escherichia coli* CRISPR-Cas3"

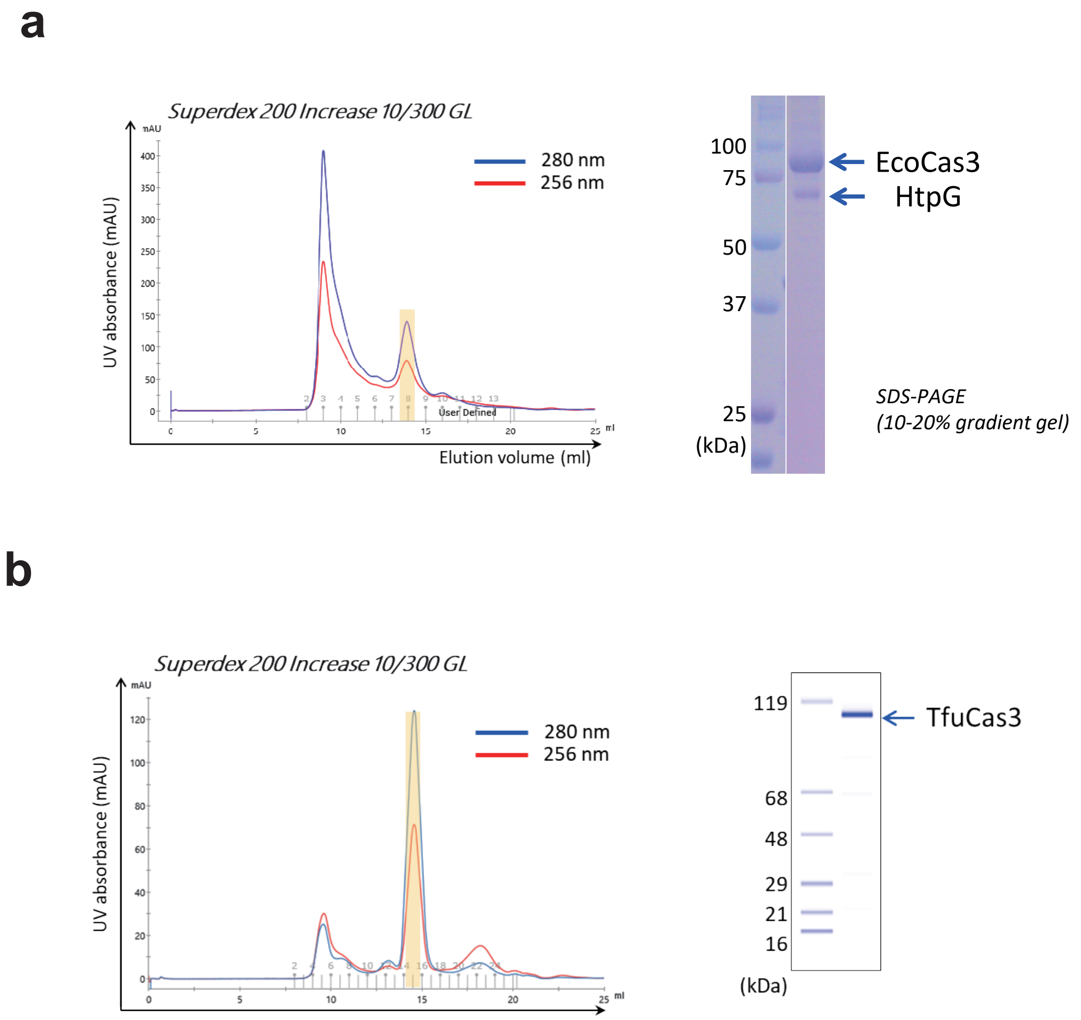


**Extended data Fig. 1. Purification of EcoCas3 and TfuCas3 recombinant proteins by the *E. coli* bacterial expression system.** (a) A limited amount of highly aggregated *E. coli* Cas3 (EcoCas3) protein was purified by size-exclusion chromatography and was evaluated by sodium dodecyl sulfate-polyacrylamide gel electrophoresis (SDS-PAGE). Co-expression of HtpG chaperon and low temperature 20°C culture were used in the *E. coli* expression system. (b) Isolation of large amounts of *Thermobifida fusca* Cas3 (TfuCas3) protein at 37°C.

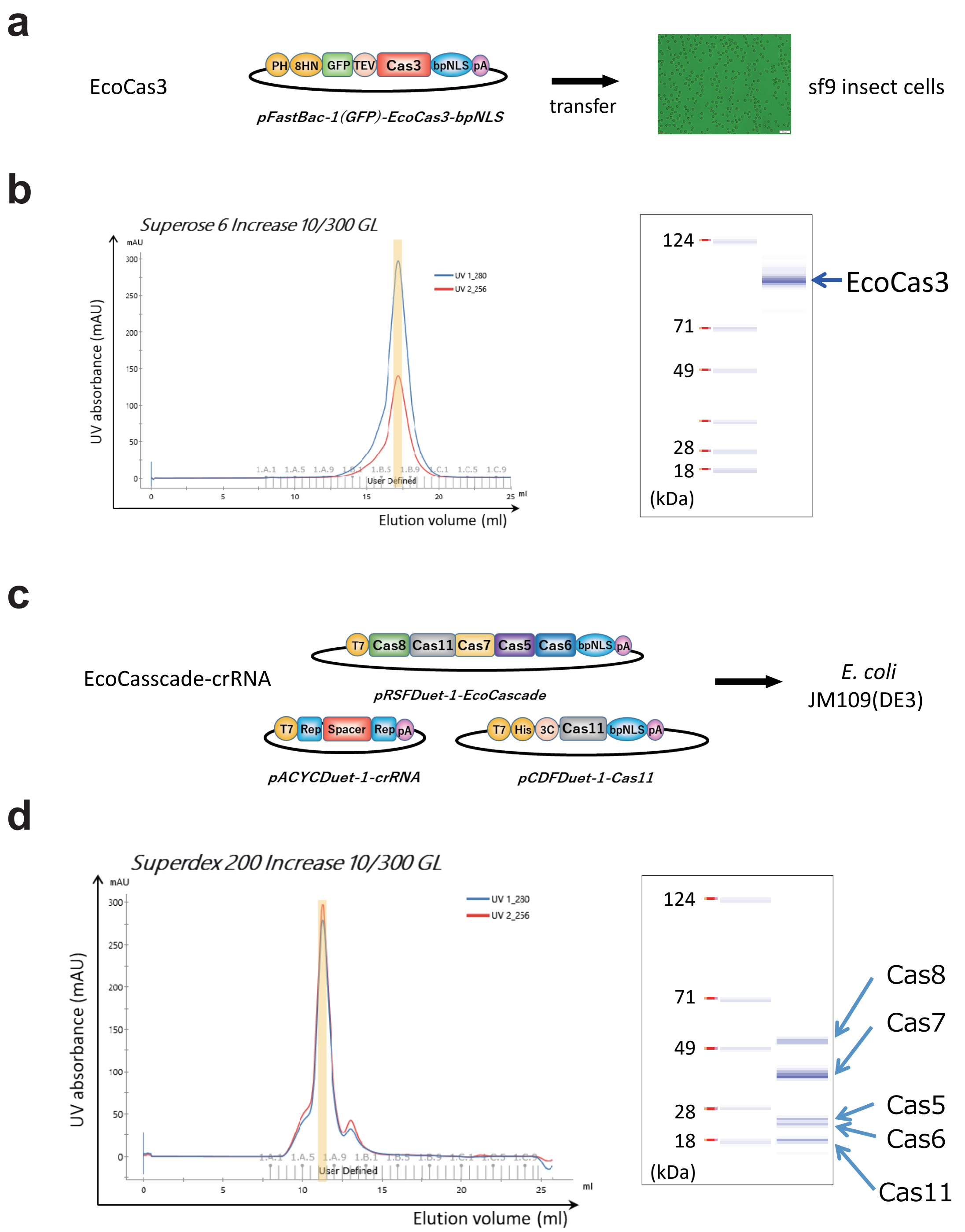


**Extended data Fig. 2. Purification of EcoCas3 and EcoCascade (Cas5, Cas6, Cas7, Cas8, and Cas11) ribonucleoproteins.** (a) A large amount of EcoCas3 protein was purified using a baculovirus expression system in Sf9 insect cells cultured at 28°C. (b) Purified EcoCas3 protein from Sf9 cells was soluble and ~95% homogeneous on SDS-PAGE. (c,d) A complex of Cas5, Cas6, Cas7, Cas8, and Cas11 proteins and crRNA was co-expressed in JM109(DE3) ***E. coli*** cultured at 37°C, purified using Ni-NTA resin, and separated by size exclusion chromatography.

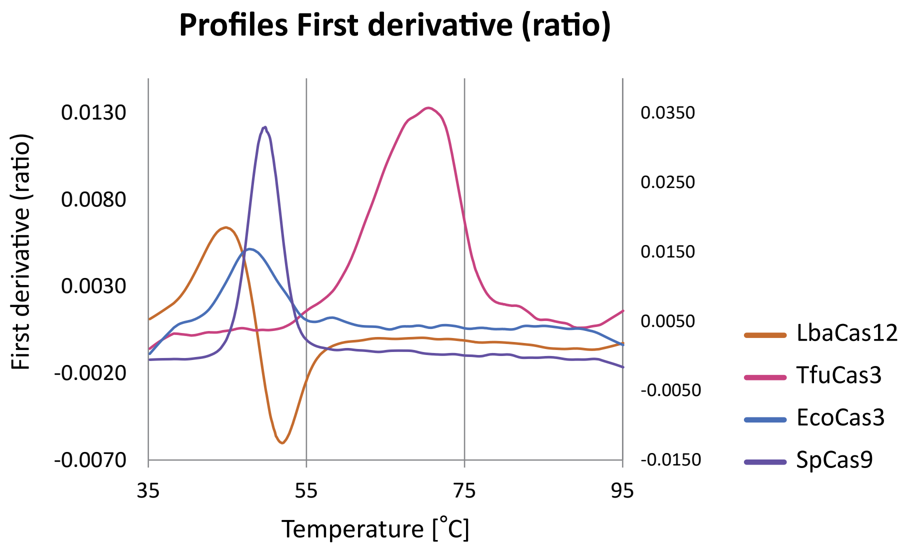


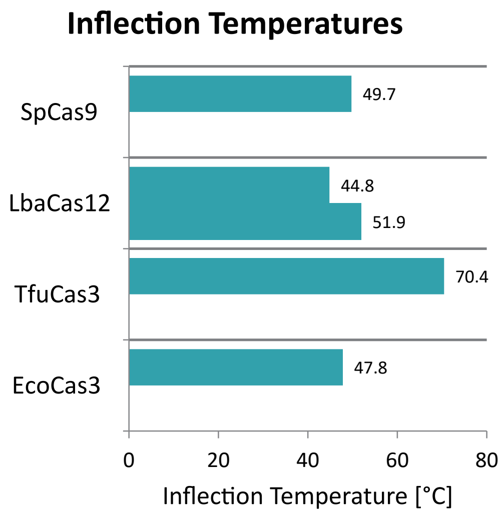


**Extended data Fig. 3. Temperature-dependent stability of recombinant EcoCas3 protein.** To evaluate the temperature-dependent stability of purified EcoCas3, we employed a modified nanoscale differential scanning fluorimetry method, nanoDSF, which determines protein stability by measuring intrinsic tryptophan or tyrosine fluorescence using the Tycho NT.6 system (NanoTemper Technologies GmbH). The profiles show the first derivative ratio of 330 and 350 nm as the temperature rises from 35°C to 95°C. EcoCas3 protein purified from Sf9 cells had a temperature inflection point (Ti) of 47.8°C, while TfuCas3 and off-the-shelf SpCas9 proteins had Ti’ s of 70.4°C and 49.7°C, respectively. LbaCas12a protein had two Tis (44.8 and 51.9°C), which may represent dimeric structural dissociation between the REC and NUC lobes.

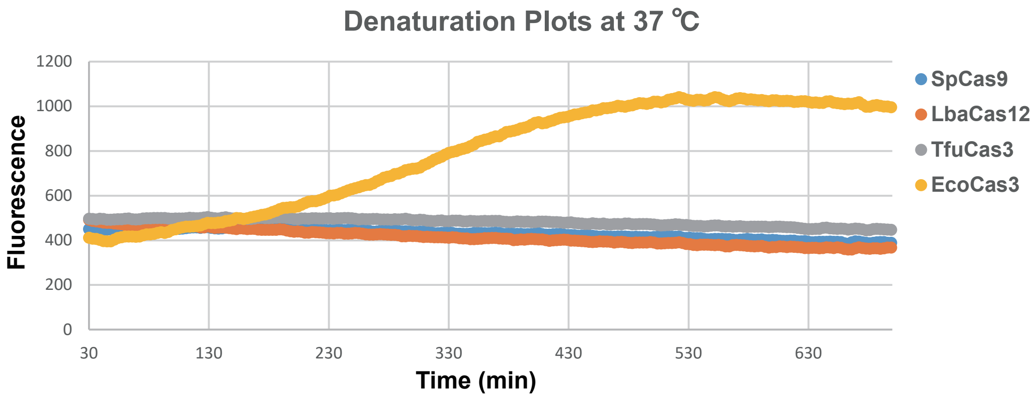


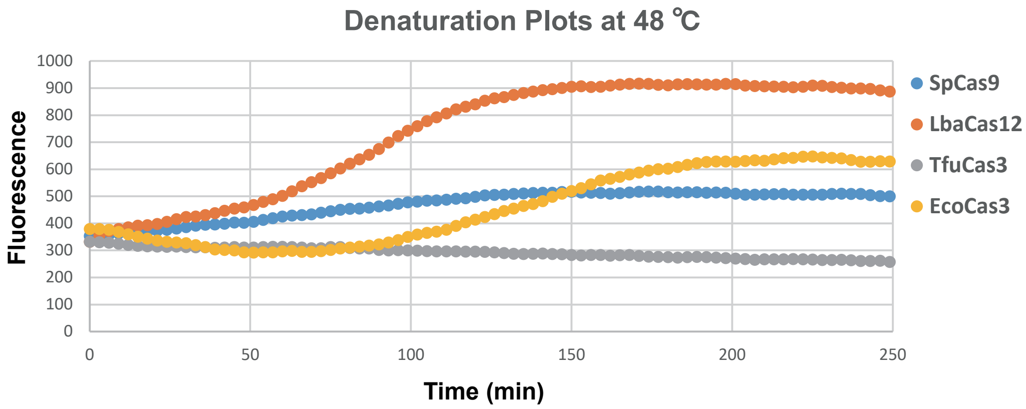


**Extended data Fig. 4. Temperature-dependent stability of recombinant EcoCas3 protein.** We evaluated temperature-dependent stability using ProteoStat detection reagent (Enzo Life Sciences), which allowed the aggregation onset temperature to be determined. Vertical and horizontal axes show fluorescence intensity, which is dependent on protein denaturation, and time, respectively. Considering thermal stability at a constant temperature of 37°C, EcoCas3 was mostly denatured in 8 h, while SpCas9, LbaCas12, and TfuCas3 were not denatured after 24 h (upper). At a constant temperature of 48°C, LbaCas12a and SpCas9 proteins showed earlier aggregation onset than EcoCas3 (lower).

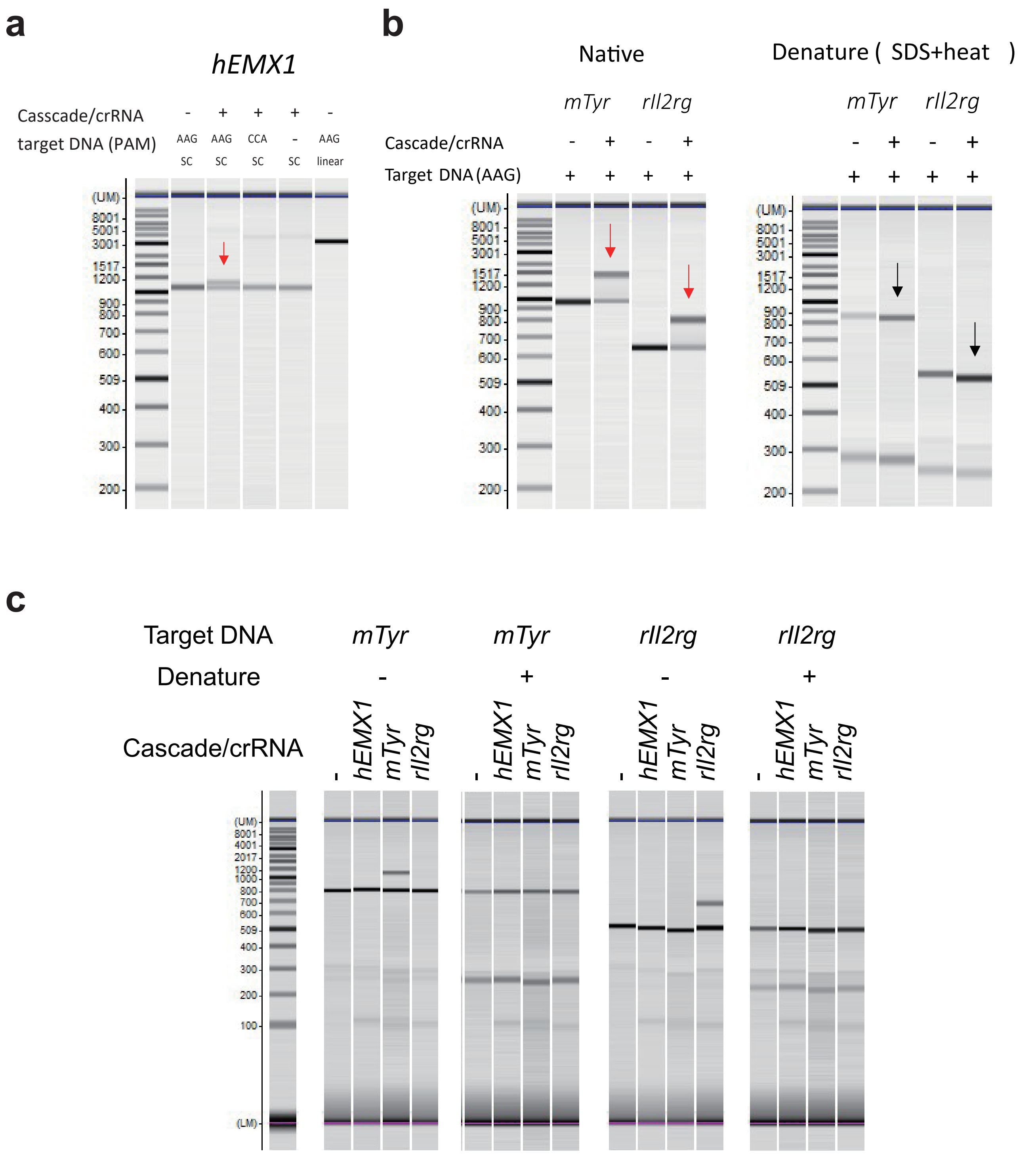


**Extended data Fig. 5. Evaluation of R-loop formation by electrophoretic mobility shift assay (EMSAs).** (a) EcoCascade/crRNA complex binds to supercoiled (SC) plasmids containing *hEMX1* spacer sequences flanked by a PAM (AAG) as shown by a red arrow, but did not bind to a nonPAM (CCA). (b) The Cascade complex binding to the PCR products of other target sequences (*mTyr* and *rIl2rg* genes). Denaturation of the Cascade complex (0.08% SDS, 95°C, 2 min) abolished DNA binding (black arrows). (c) Exchange of nucleotide pairs between crRNAs and target sequences abolished the binding, indicating the specificity of target recognition by the EcoCascade/crRNA complex.

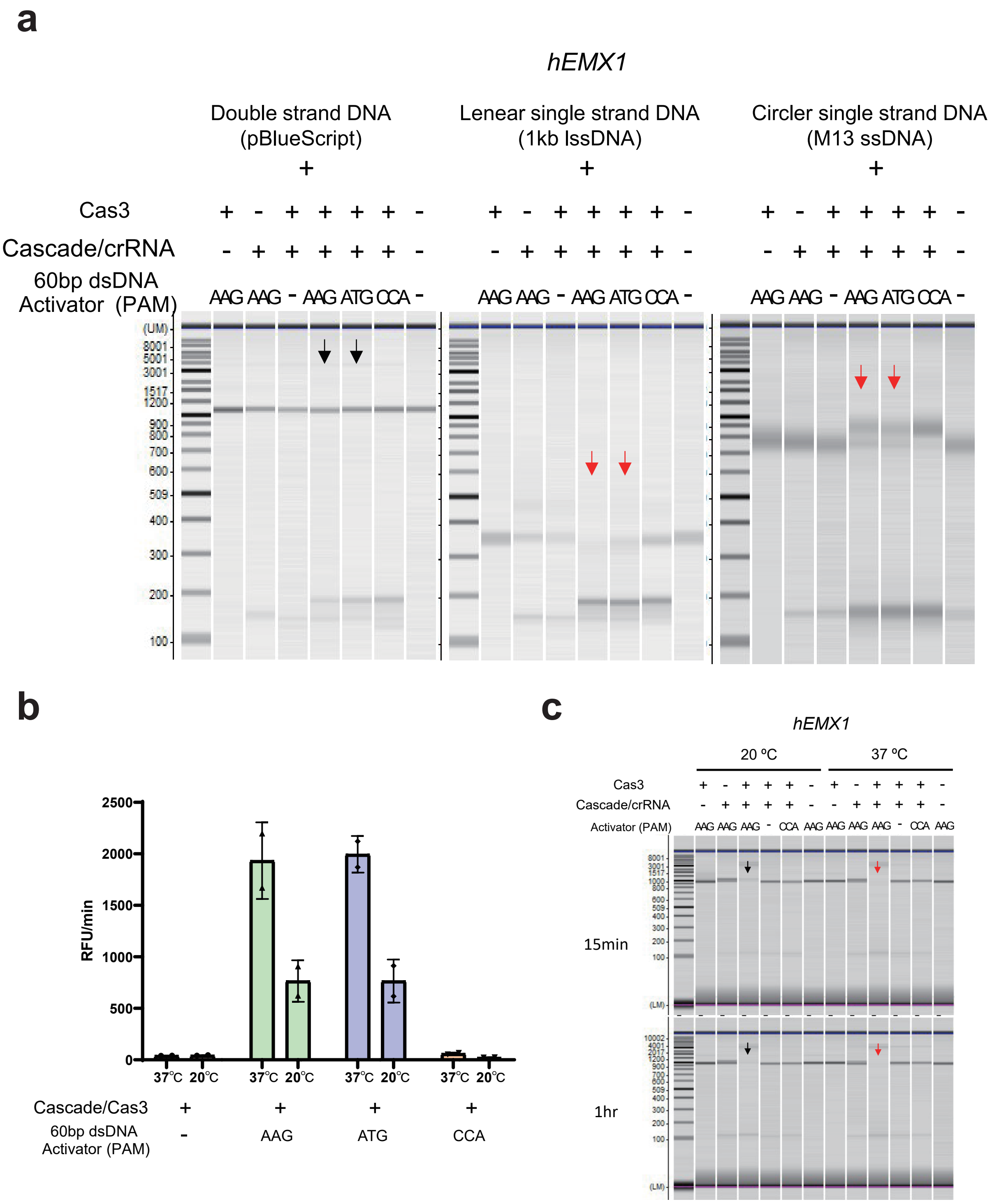


**Extended data Fig. 6. Collateral cleavages of nearby non-specific ssDNAs.** (a) EcoCas3 possesses collateral non-specific single-stranded DNA (ssDNA) cleavage activity after target-specific double-stranded DNA (dsDNA) cleavage. Circular ssDNA M13 phage and linearized long ssDNA were degraded after incubation for 1 h at 37°C (red arrows), but circular pBlueScript dsDNA was not cleaved (black arrows). (b) Comparison of collateral cleavage activity between 37°C and 20°C. (c) *In vitro* reconstitution assay for target DNA degradation also showed higher activity at 37°C (red arrows) than at 20°C (black arrows).

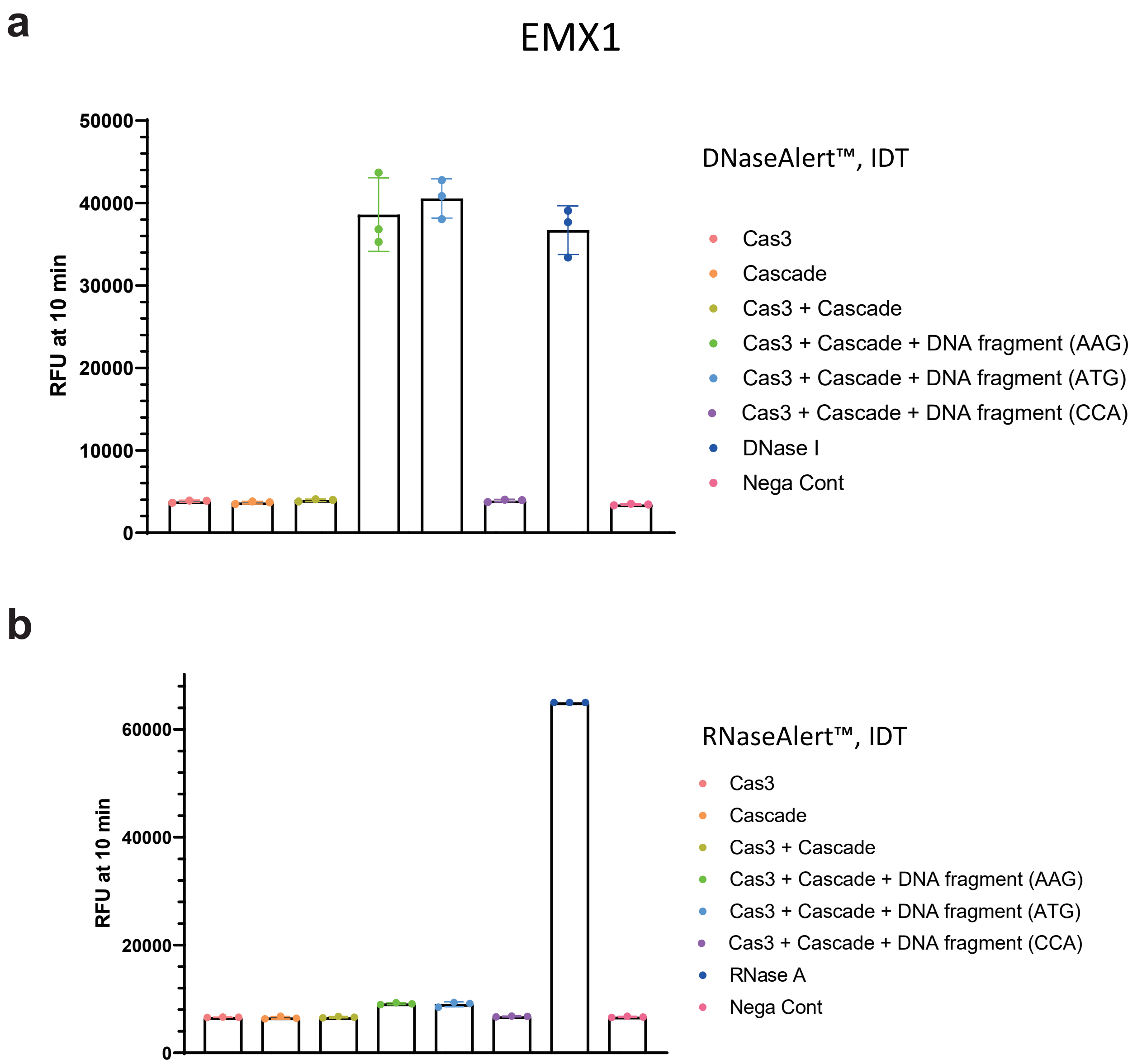


**Extended data Fig. 7. Collateral cleavage activity for ssDNAs and ssRNAs.** (a) Using fluorescent reporter DNA oligonucleotides (DNaseAlert™, IDT), we detected collateral ssDNA cleavage activity by assembling EcoCas3, EcoCascade RNPs and dsDNA fragments that included target sequences (*hEMX1* or *mTyr*) flanked by PAM-AAG or -ATG, but not with PAM-CCA. (b) Little or no RNase activity was detected with fluorescent reporter RNA oligonucleotides (RNaseAlert™, IDT) by assembling EcoCas3, EcoCascade RNPs and dsDNA fragments.

a

b

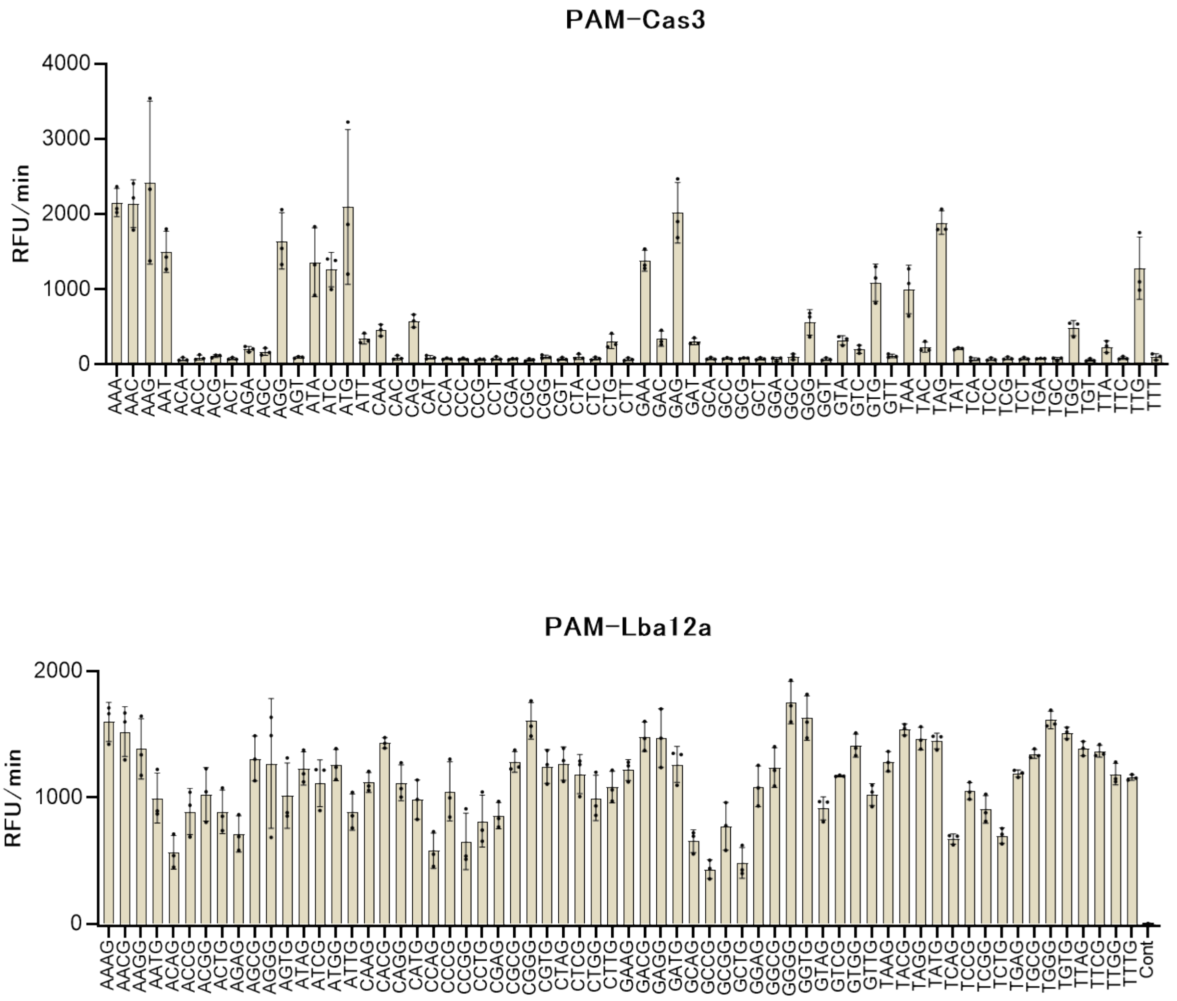


**Extended data Fig. 8. Specific PAM recognition for collateral ssDNA cleavage by EcoCas3.** (a,b) Screening of all 64 possible target sites containing each of the three-nucleotide PAM sequences for *trans* cleavage activity by EcoCas3 (a) and LbaCas12a (b).

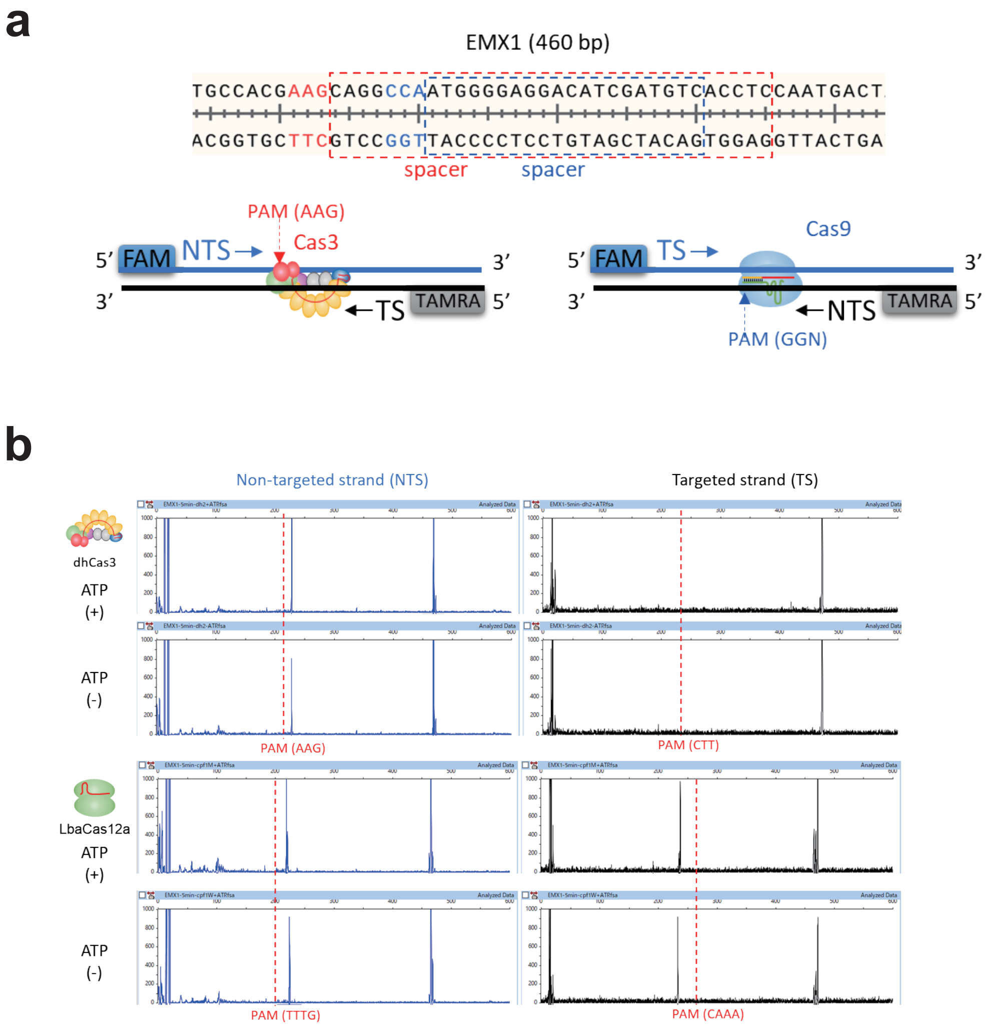


**Extended data Fig. 9. The dsDNA cleavage assay.** (a) Schematic depictions of the dsDNA cleavage assay. Fluorescently-labeled target dsDNA substrates, 5′-NTS-FAM and 5′-TS-TAMRA, to visualize dsDNA cleavage. (b) The dhCas3 SF2 domain mutant cleaves the NTS in *cis*, but not the TS in *trans* in ATP (+ and -) reaction buffer, while LbaCas12a cleaves both the NTS and TS in ATP (+ and -) reaction buffer.

**a**

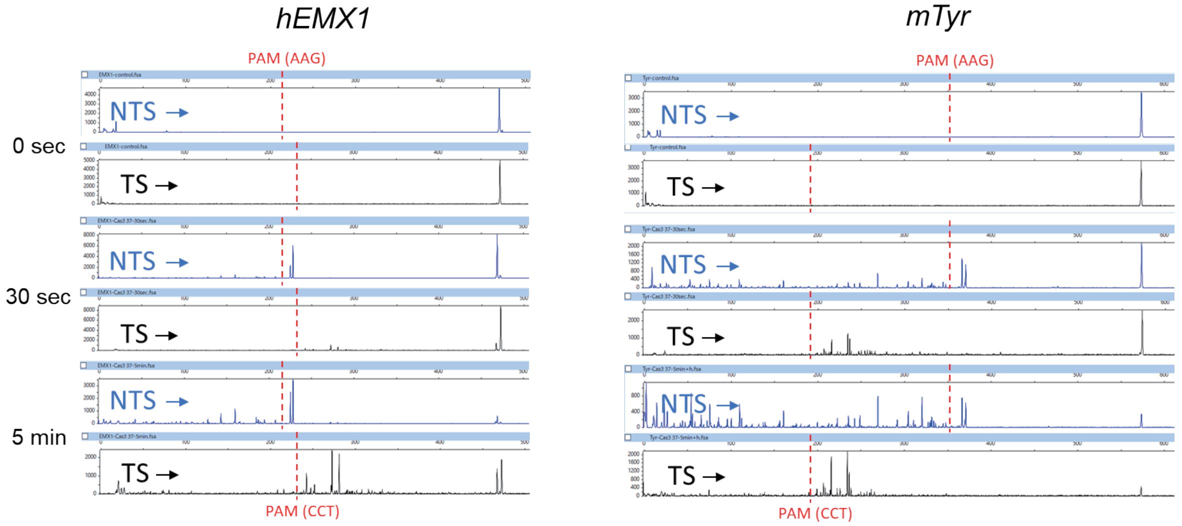


**b**

| Sample File Name | Dye | Sample Peak | Size | Height |
|---|---|---|---|---|
| EMX1-Cas3 37-30sec.fsa | B | 29 | 143.2 | 482 |
| EMX1-Cas3 37-30sec.fsa | B | 31 | 159.7 | 661 |
| EMX1-Cas3 37-30sec.fsa | B | 42 | 224.9 | 2391 |
| EMX1-Cas3 37-30sec.fsa | B | 43 | 228.0 | 6101 |
| EMX1-Cas3 37-30sec.fsa | B | 45 | 468.8 | 8146 |
| EMX1-Cas3 37-30sec.fsa | Y | 67 | 272.3 | 1103 |
| EMX1-Cas3 37-30sec.fsa | Y | 70 | 280.6 | 700 |
| EMX1-Cas3 37-30sec.fsa | Y | 130 | 467.0 | 1464 |
| EMX1-Cas3 37-30sec.fsa | Y | 133 | 472.3 | 8801 |
| EMX1-Cas3 37-5min.fsa | B | 33 | 143.1 | 810 |
| EMX1-Cas3 37-5min.fsa | B | 35 | 159.7 | 1172 |
| EMX1-Cas3 37-5min.fsa | B | 39 | 184.8 | 519 |
| EMX1-Cas3 37-5min.fsa | B | 51 | 225.1 | 2510 |
| EMX1-Cas3 37-5min.fsa | B | 52 | 228.1 | 3525 |
| EMX1-Cas3 37-5min.fsa | B | 57 | 468.9 | 619 |
| EMX1-Cas3 37-5min.fsa | Y | 16 | 20.5 | 728 |
| EMX1-Cas3 37-5min.fsa | Y | 87 | 242.2 | 1145 |
| EMX1-Cas3 37-5min.fsa | Y | 92 | 251.6 | 527 |
| EMX1-Cas3 37-5min.fsa | Y | 101 | 271.3 | 491 |
| EMX1-Cas3 37-5min.fsa | Y | 102 | 272.4 | 2415 |
| EMX1-Cas3 37-5min.fsa | Y | 106 | 280.8 | 2203 |
| EMX1-Cas3 37-5min.fsa | Y | 167 | 467.0 | 1393 |
| EMX1-Cas3 37-5min.fsa | Y | 170 | 472.4 | 1901 |
| EMX1-Cas9 37-5min.fsa | B | 6 | 229.2 | 15251 |
| EMX1-Cas9 37-5min.fsa | B | 9 | 468.9 | 10292 |
| EMX1-Cas9 37-5min.fsa | Y | 55 | 229.0 | 593 |
| EMX1-Cas9 37-5min.fsa | Y | 57 | 236.2 | 583 |
| EMX1-Cas9 37-5min.fsa | Y | 58 | 237.0 | 1500 |
| EMX1-Cas9 37-5min.fsa | Y | 59 | 238.0 | 7933 |
| EMX1-Cas9 37-5min.fsa | Y | 109 | 468.8 | 472 |
| EMX1-Cas9 37-5min.fsa | Y | 110 | 472.4 | 9178 |
| EMX1-control.fsa | B | 8 | 469.0 | 4667 |
| EMX1-control.fsa | Y | 118 | 472.5 | 5050 |
| Tyr-Cas3 37-30sec.fsa | B | 16 | 52.5 | 401 |
| Tyr-Cas3 37-30sec.fsa | B | 22 | 75.0 | 401 |
| Tyr-Cas3 37-30sec.fsa | B | 28 | 109.4 | 406 |
| Tyr-Cas3 37-30sec.fsa | B | 56 | 269.1 | 699 |
| Tyr-Cas3 37-30sec.fsa | B | 63 | 320.2 | 487 |
| Tyr-Cas3 37-30sec.fsa | B | 75 | 366.3 | 1410 |
| Tyr-Cas3 37-30sec.fsa | B | 77 | 370.5 | 1148 |
| Tyr-Cas3 37-30sec.fsa | B | 79 | 573.3 | 2171 |
| Tyr-Cas3 37-30sec.fsa | Y | 83 | 215.7 | 914 |
| Tyr-Cas3 37-30sec.fsa | Y | 91 | 234.2 | 1258 |
| Tyr-Cas3 37-30sec.fsa | Y | 204 | 573.2 | 2608 |
| Tyr-Cas3 37-5min.fsa | B | 14 | 53.5 | 610 |
| Tyr-Cas3 37-5min.fsa | B | 20 | 74.4 | 447 |
| Tyr-Cas3 37-5min.fsa | B | 21 | 75.2 | 544 |
| Tyr-Cas3 37-5min.fsa | B | 29 | 109.4 | 536 |
| Tyr-Cas3 37-5min.fsa | B | 53 | 269.0 | 585 |
| Tyr-Cas3 37-5min.fsa | B | 61 | 320.2 | 436 |
| Tyr-Cas3 37-5min.fsa | B | 70 | 366.3 | 456 |
| Tyr-Cas3 37-5min.fsa | Y | 85 | 215.6 | 1099 |
| Tyr-Cas3 37-5min.fsa | Y | 93 | 234.3 | 987 |
| Tyr-Cas3 37-5min.fsa | Y | 94 | 236.4 | 514 |
| Tyr-Cas3 37-5min+h.fsa | B | 8 | 22.6 | 609 |
| Tyr-Cas3 37-5min+h.fsa | B | 9 | 25.9 | 408 |
| Tyr-Cas3 37-5min+h.fsa | B | 16 | 53.6 | 873 |
| Tyr-Cas3 37-5min+h.fsa | B | 23 | 75.1 | 594 |
| Tyr-Cas3 37-5min+h.fsa | B | 31 | 109.5 | 592 |
| Tyr-Cas3 37-5min+h.fsa | B | 41 | 160.4 | 425 |
| Tyr-Cas3 37-5min+h.fsa | B | 60 | 269.1 | 806 |
| Tyr-Cas3 37-5min+h.fsa | B | 65 | 304.2 | 424 |
| Tyr-Cas3 37-5min+h.fsa | B | 70 | 320.1 | 780 |
| Tyr-Cas3 37-5min+h.fsa | B | 82 | 366.3 | 754 |
| Tyr-Cas3 37-5min+h.fsa | B | 84 | 370.5 | 631 |
| Tyr-Cas3 37-5min+h.fsa | Y | 78 | 206.7 | 624 |
| Tyr-Cas3 37-5min+h.fsa | Y | 79 | 208.8 | 403 |
| Tyr-Cas3 37-5min+h.fsa | Y | 84 | 215.6 | 1898 |
| Tyr-Cas3 37-5min+h.fsa | Y | 94 | 234.3 | 2154 |
| Tyr-Cas3 37-5min+h.fsa | Y | 95 | 236.4 | 1147 |
| Tyr-Cas3 37-5min+h.fsa | Y | 96 | 238.4 | 417 |
| Tyr-Cas3 37-5min+h.fsa | Y | 183 | 573.1 | 486 |
| Tyr-control.fsa | B | 7 | 573.3 | 3493 |
| Tyr-control.fsa | Y | 148 | 573.2 | 3565 |

**Extended data Fig. 10. The dsDNA cleavage assay.** (a) Comparison of 30 sec (short) and 5 min (long) incubation for the dsDNA cleavage assay (*hEMX1* or *mTyr*). (b) The size and patterns of cleaved fragments after short and long incubation in the dsDNA cleavage assay.

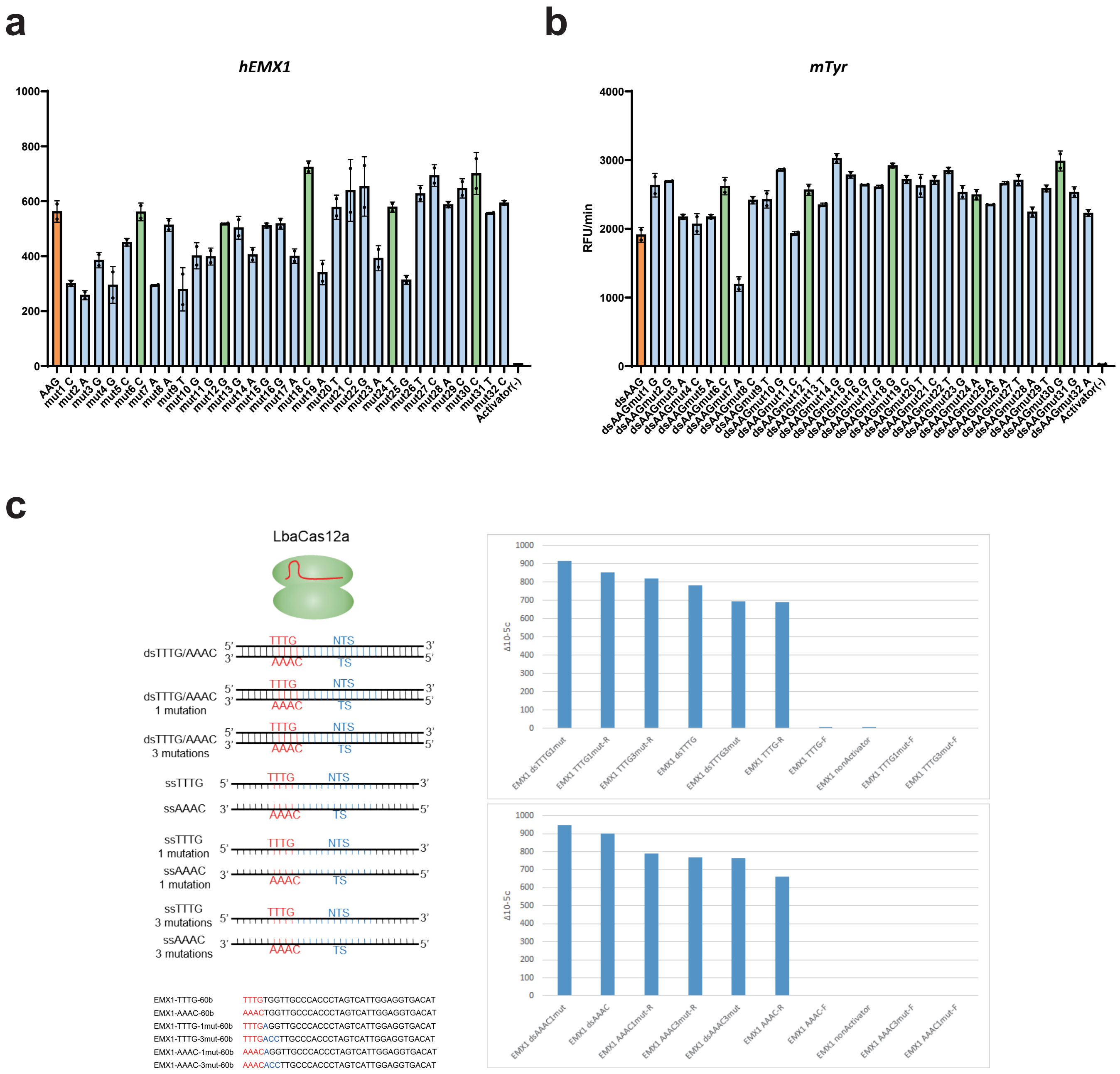


**Extended data Fig. 11. The effect of mismatch of each 32-nt spacer sequence on collateral cleavage activity.** (a,b) A single mismatch in the spacer region has a little or no effect on collateral cleavage activity of *hEMX1* target (a) and *mTyr* target (b). (c) One to three mismatches in the PAM sites and the spacer region have little effect on collateral cleavage activity by LbaCas12a.

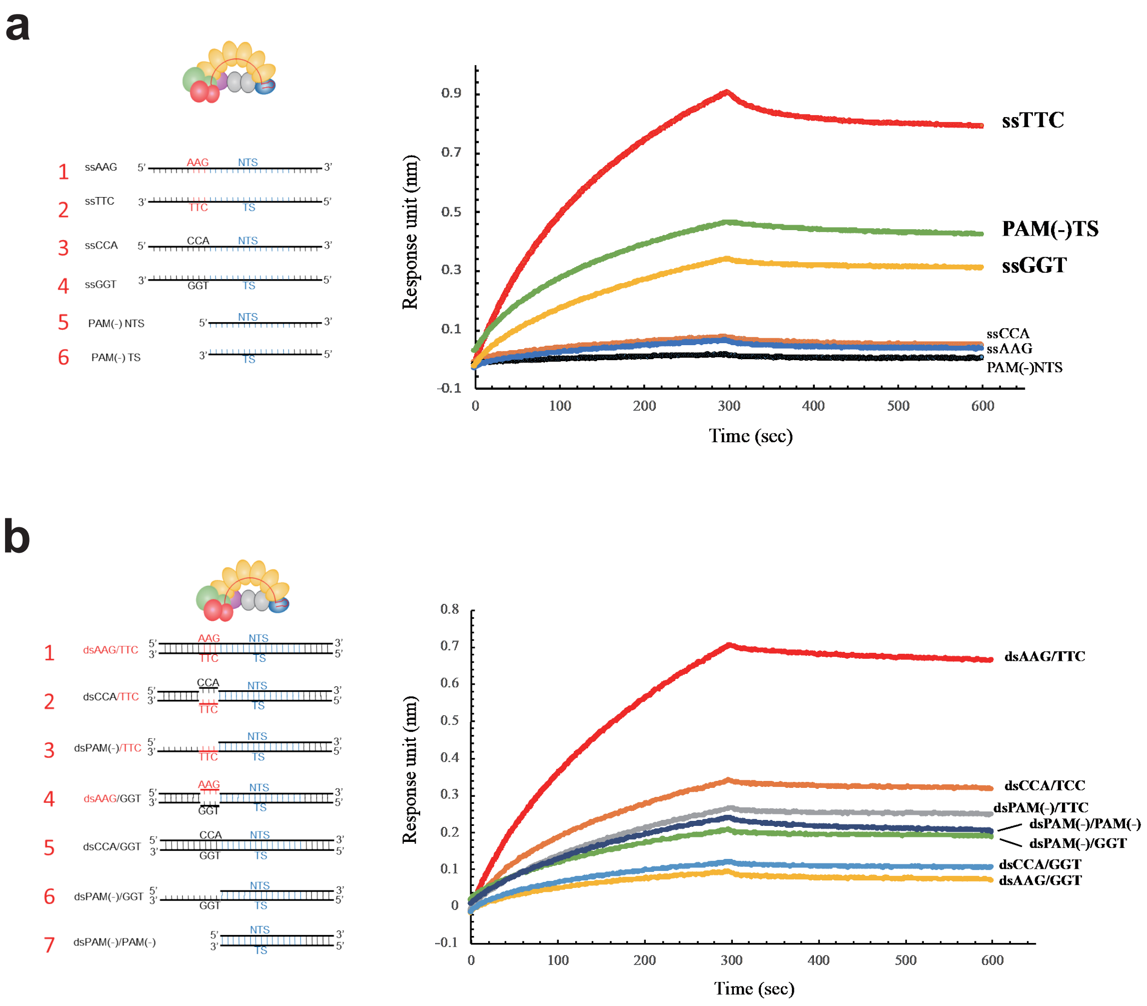


**Extended data Fig. 12. EcoCascade-target DNA associations and dissociations measured by the Octet RED 96 System.** (a) crRNA-complementary ssDNA with TS-PAM (TTC) showing collateral cleavage (Fig. 2c) represents higher association than TS-nonPAM or TS-PAMless. (b) The interactions between Cascade and dsDNAs containing paired PAM (AAG-TTC), unpaired PAM between TS-PAM (TTC) and NTS-nonPAM (CCA), and unpaired PAM between NTS-PAM (AAG) and TS-nonPAM (GGT), correspond with the results from the collateral cleavage assay (Fig. 2f).

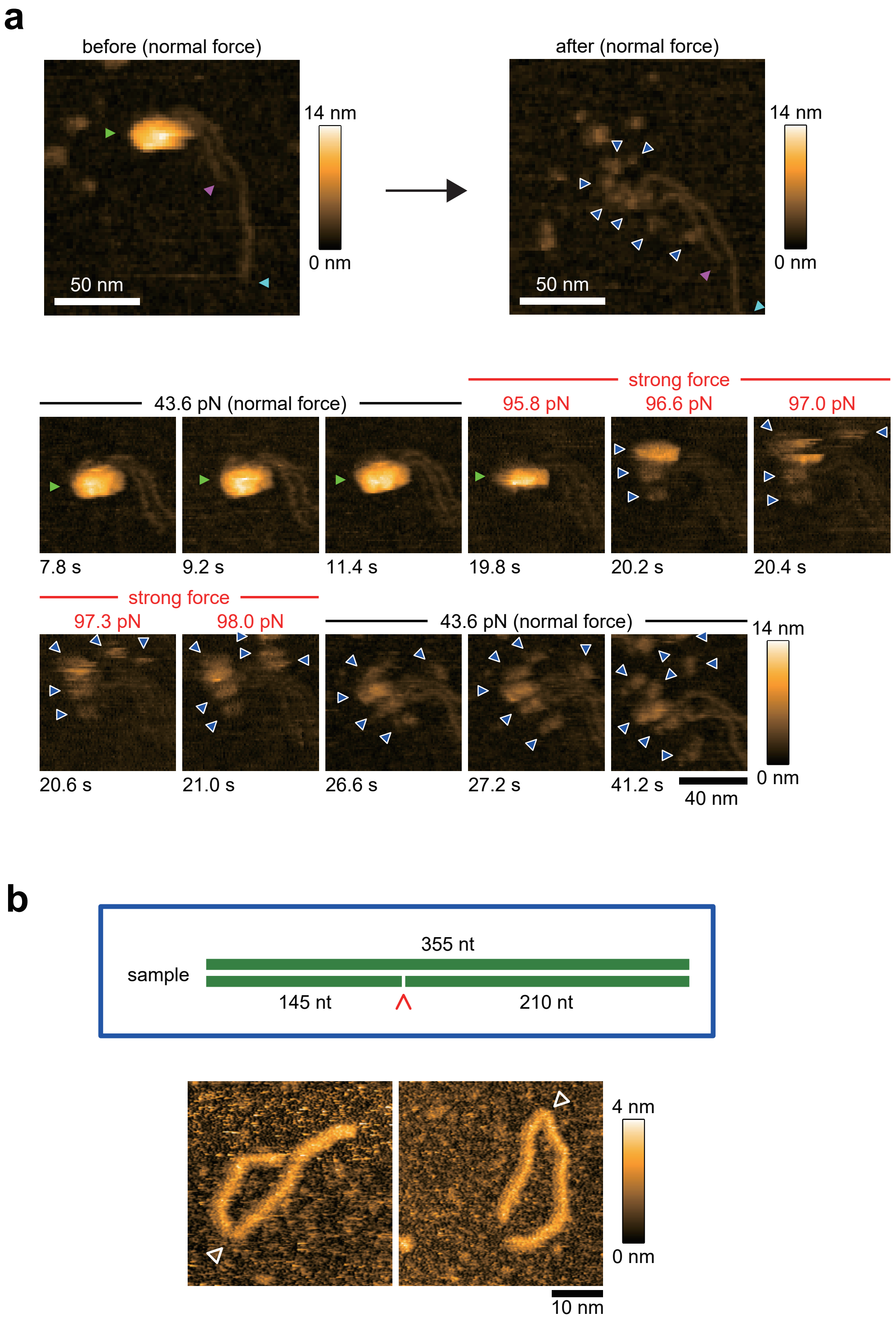


**Extended data Fig. 13. Dynamic visualization of CRISPR interference by hs-AFM.** (a) By applying excessive force, the EcoCascade RNP body (green triangle) was broken and separated into multiple Cas effector components (white triangle) (video 3). (b) The shape of bent DNA (white arrows) resulting from artificial nicking by Nb.BsrDI endonuclease.

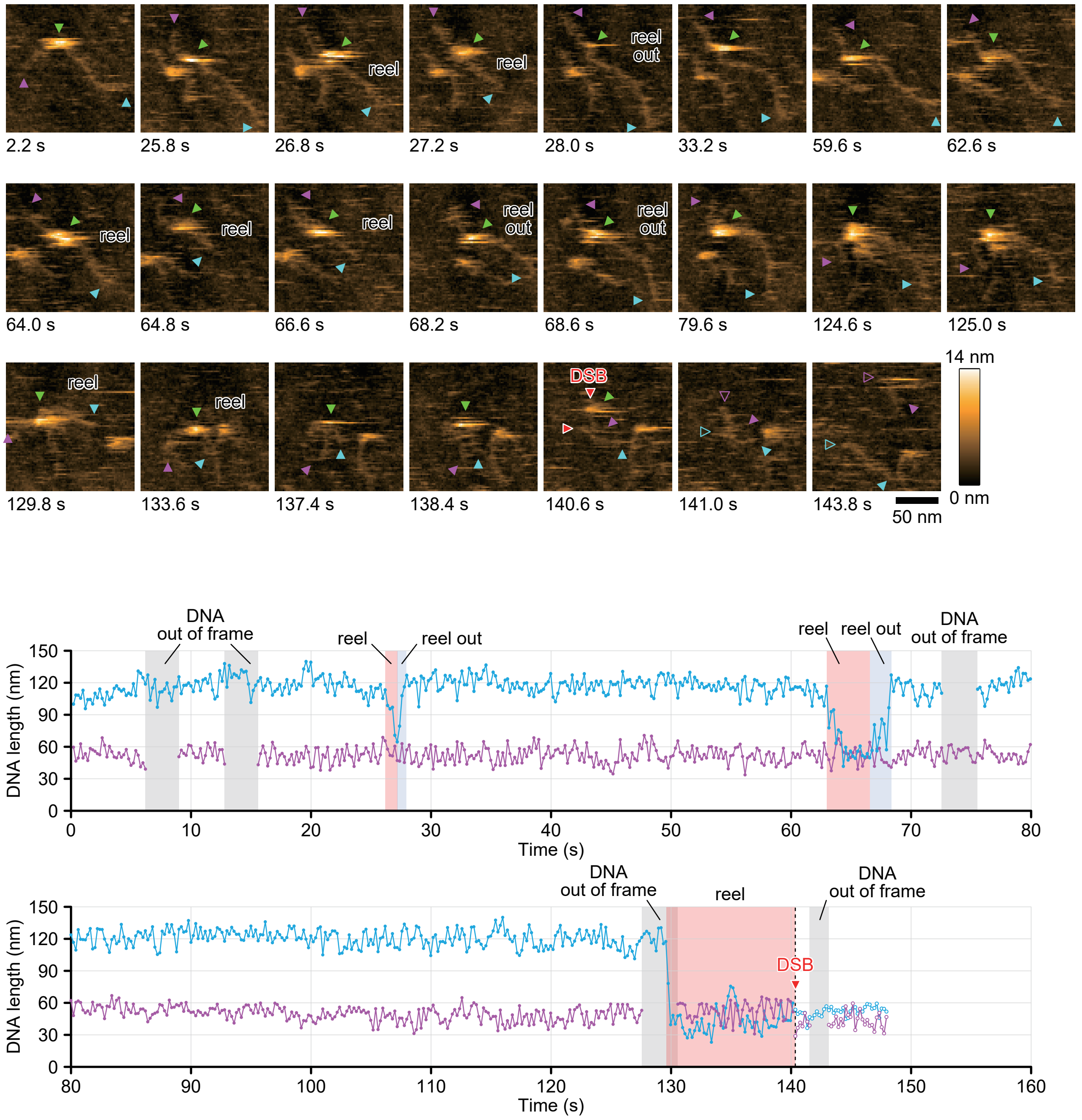


**Extended data Fig. 14. Dynamic visualization of CRISPR interference by hs-AFM.** In ATP (+) reaction buffer, the EcoCas3-Cascade complex repeatedly reels and releases the longer side of the DNA (blue arrows) and then cleaves it with a DSB (red arrows) (video 5).
